# *Ex vivo* and *in vivo* HIV-1 latency reversal by “Mukungulu,” a protein kinase C-activating African medicinal plant extract

**DOI:** 10.1101/2024.09.15.613141

**Authors:** Khumoekae Richard, Zhe Yuan, Hsin-Yao Tang, Aaron R. Goldman, Riza Kuthu, Boingotlo Raphane, Emery T. Register, Paridhima Sharma, Brian N. Ross, Jessicamarie Morris, David E. Williams, Carol Cheney, Guoxin Wu, Karam Mounzer, Gregory M. Laird, Paul Zuck, Raymond J. Andersen, Sundana Simonambango, Kerstin Andrae-Marobela, Ian Tietjen, Luis J. Montaner

**Affiliations:** The Wistar Institute, Philadelphia, PA, USA; University of Botswana, Gaborone, Botswana; Departments of Chemistry and Earth, Ocean & Atmospheric Sciences, University of British Columbia, Vancouver, BC V6T 1Z4, Canada; Merck & Co., Inc., Rahway, NJ, USA; Jonathan Lax Immune Disorders Treatment Center, Philadelphia Field Initiating Group for HIV-1 Trials, Philadelphia, PA, USA; AccelevirDx, Baltimore, MD, USA; Kwame (Legwame) Traditional Association, P.O. Box 3481, Mmadinare, Botswana

## Abstract

Current HIV latency reversing agents (LRAs) have had limited success in clinic, indicating the need for new strategies that can reactivate and/or eliminate HIV reservoirs. “Mukungulu,” prepared from the bark of *Croton megalobotrys* Müll. Arg., is traditionally used for HIV/AIDS management in Northern Botswana despite containing an abundance of protein kinase C (PKC)-activating phorbol esters (“namushens”). Here we show that Mukungulu is tolerated in mice at up to 12.5 mg/kg while potently reversing latency in antiretroviral therapy (ART)-suppressed HIV-infected humanized mice at 5 mg/kg. In peripheral blood mononuclear cells (PBMC) and isolated CD4+ T-cells from ART-suppressed people living with HIV-1, 1 µg/mL Mukungulu reverses latency on par with or superior to anti-CD3/CD28 positive control, as measured by HIV gag-p24 protein expression, where the magnitude of HIV reactivation in PBMC corresponds to intact proviral burden levels in CD4+ T-cells. Bioassay-guided fractionation identifies 5 namushen phorbol ester compounds that reactivate HIV expression, yet namushens alone do not match Mukungulu’s activity, suggesting additional enhancing factors. Together, these results identify Mukungulu as a robust natural LRA which may warrant inclusion in future LRA-based HIV cure and ART-free remission efforts.

## Introduction

While antiretroviral therapy (ART) is a remarkable milestone that has reduced HIV/AIDS-related morbidities and mortalities globally, ART is not curative. A major obstacle to HIV eradication is the presence of inducible, replication-competent proviral DNA that persists in cellular reservoirs, particularly in CD4+ T-cells, which can reactivate at any time to produce infectious virus ^1–2^. Because of this, people living with HIV (PLWH) must remain on ART for life. Long-term use of ART is also increasingly linked to non-AIDS co-morbidities including renal disorders, cancer, and cardiovascular diseases, potentially due to residual viral antigen production, ongoing immune activation, and/or chronic inflammation ^3–5^. As a result, new therapies that can support lowering viral burden on ART by targeting viral reservoirs continue to be needed.

One important strategy to identify and eliminate HIV reservoirs involves the use of latency reversing agents (LRAs) that activate HIV-1 production (in the co-presence of ART to eliminate reservoir re-seeding) to expose latently infected cells to immune-mediated clearance, an approach colloquially known as “shock-and-kill” or “kick-and-kill” ^6–8^. Numerous LRAs have been identified encompassing different molecular mechanisms of action such as histone deacetylase inhibitors (HDACis), activators of protein kinase C (PKC) signaling, bromodomain and extra-terminal bromodomain inhibitors, DNA methyltransferase inhibitors, histone methyltransferase inhibitors, programmed cell death protein-1 inhibitors, toll-like receptor agonists, and noncanonical NF-κB agonists, among others ^8^. However, most of these LRAs remain poorly characterized in both *ex vivo* models using primary blood cells from PLWH and/or *in vivo* animal models ^9–10,29^. Furthermore, LRAs that have been tested to date in humans such as HDACis largely represent repurposed anti-cancer drugs with limited to no impact on viral reservoir size in PLWH ^9–10,29^. By contrast, in spite of greater activity *in vitro*, the large majority of PKC activators have not advanced beyond *ex vivo* models due to the risk that single PKC activator compounds, for example phorbol esters like phorbol 12-myristate 13-acetate (PMA) or prostratin, may cause widespread T cell activation and *in vivo* toxicity ^11–12^.

Using a reverse pharmacology approach ^13^, medicinal plants have been documented that are traditionally used for HIV/AIDS management in Sub-Saharan Africa and elsewhere, including some with latency-reversing properties ^14–15^. For example, we previously described the traditional use of *Croton megalobotrys* Müll Arg. bark preparations, locally called “Mukungulu,” to manage HIV/AIDS in Northern Botswana ^14,16^. Though Botswana introduced free and universal ART access in 2002, many PLWH initially hesitated to enroll in these programs due to perceived stigma and discrimination and instead chose to rely on local primary health care providers, which included traditional healers. We documented that Mukungulu is part of a three-step traditional treatment regimen which was offered to patients in Northern Botswana during that time and which is now also used as a supplement to standard ART ^16^. According to the healers, patients are treated if presenting with substantial weight loss, chronic diarrhea, fevers, skin infections or wounds, and/or lethargy. The first two steps of the treatment regimen consist of separate plant preparations (from *Cassia siberiana* D.C. roots and *Vitex doniana* (Sweet) bark) taken over several weeks which also inhibit HIV replication *in vitro* ^16^. Mukungulu is then administered as a single dose (third step in regimen), where the patient remains under close observation by the healer for 48-72 hours. After recovery, patients are instructed to continue use of ART, obtain plasma viral load results from a local clinic after 3 weeks, and not repeat Mukungulu treatment for at least 6-12 months ^14,16^. Following treatment, patients report improved weight gain, recovery from chronic diarrhea, fewer fevers, wound healing, and increased energy for at least 3 months ^14,16^. Notably, we found that crude Mukungulu extract reversed HIV latency in cell lines and contained phorbol esters (namushens 1 and 2) that structurally resemble PMA and prostratin and were sensitive to PKC inhibition *in vitro*, supporting a mode of action involving PKC activation ^14^. These properties, together with Mukungulu’s traditional single dose following anticipated viral suppression, suggested it may function as a novel LRA.

To further investigate the potential of Mukungulu as an LRA acting through PKC activation, we describe here the latency reversing properties of Mukungulu and its active components in total PBMC and isolated CD4+ T-cells from ART-suppressed PLWH. We also document Mukungulu’s activity when used in ART-suppressed HIV-infected humanized mice, where we observe robust latency reversal as well as *in vivo* tolerability.

## Results

### Isolation of phorbol ester-class LRAs (“namushens”) from crude Mukungulu extract

We previously isolated two phorbol esters (namushen 1 and 2, named for the healer who first communicated traditional use of Mukungulu) from Mukungulu that could reverse HIV latency *in vitro* ^14,16^. We therefore began by expanding the characterization of Mukungulu extract to identify if any other active components apart from namushens 1 and 2 were present. To do this, we obtained 81.3 grams of an oily dark brown CH_2_Cl_2_/methanol crude extract from a collection of *Croton megalobotrys* bark powder and subjected it to bioassay-guided fractionation using J-Lat 9.2 cells and using previously-described *in vitro* approaches ^14,17^ (see **Supplementary Materials**). Briefly, J-Lat 9.2 cells were treated with compounds for 24 hours, and GFP reporter expression, a marker of provirus expression, ^18^ was assessed in live cells by flow cytometry. This approach led to re-isolation of namushens 1 and 2, as expected, but also three new phorbol ester species which we named namushens 3, 4 and 5 (**Fig. 1A**, **Table S1**, **Fig. S1-S5**). Notably, no bioactive fractions were identified that lacked namushens, establishing that phorbol esters are the primary driver of *in vitro* HIV latency reversal.

**Fig. 1.**
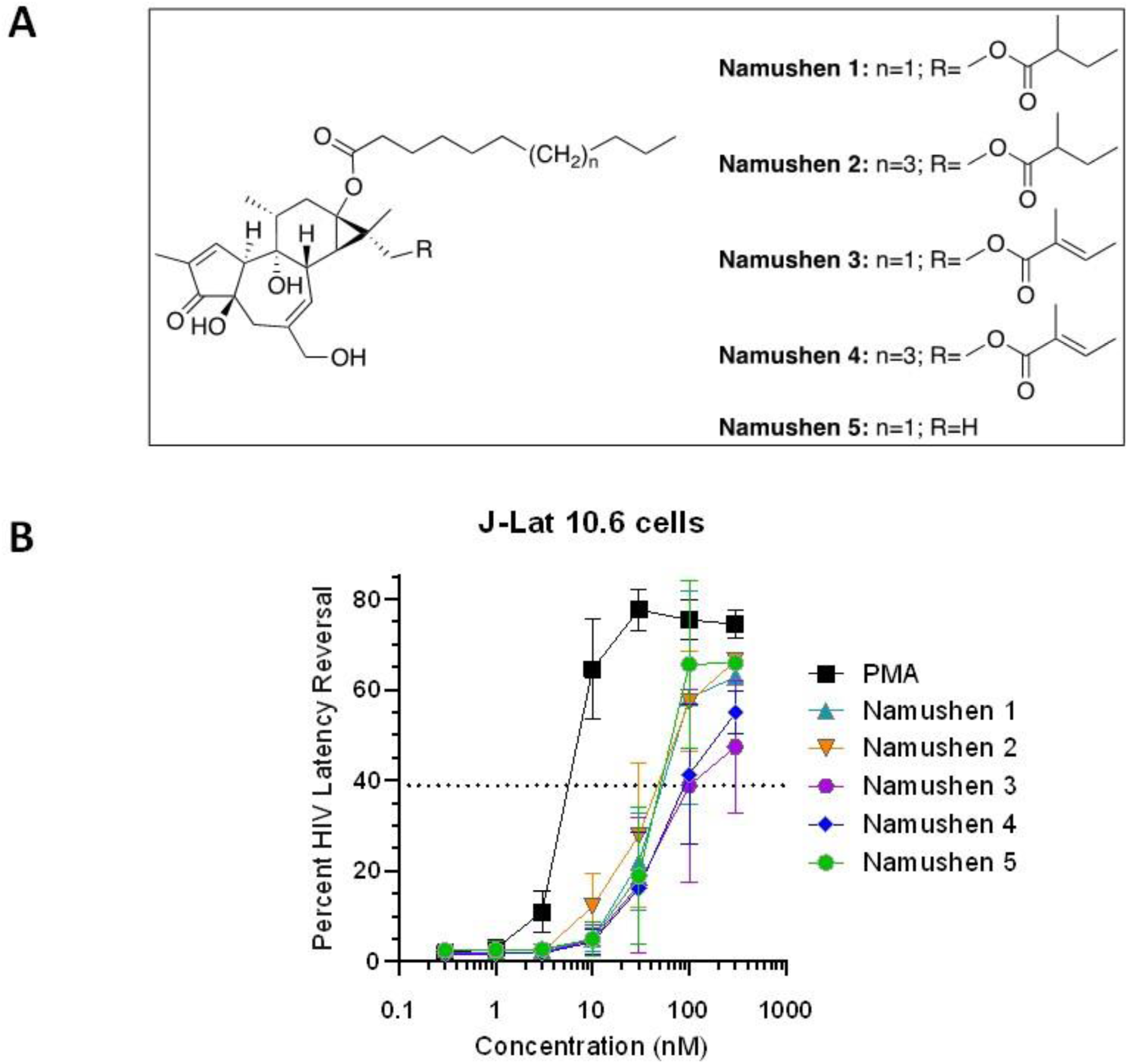
Isolation of namushen phorbol esters as major LRA-components of crude Mukungulu extract. **A,** Structures of 5 isolated namushen phorbol esters. **B,** Effects of namushens and control phorbol ester PMA on HIV latency reversal in J-Lat 10.6 cells.

To reconfirm HIV latency reversal by purified namushens, we assessed their activity as compared to PMA in J-Lat 10.6 cells, which resemble J-Lat 9.2 cells except for a different genomic location of provirus integration. With this approach, we determined a half-maximal effective concentration (EC_50_) of 4.1 nM for PMA compared to EC_50_s of 48.0, 41.2, 135.2, 99.5, and 40.9 nM for namushens 1-5, respectively (**Fig. 1B**), indicating that namushens have roughly 10 to 30-fold lower activity than PMA and that the distal double bond of the hydrophobic tail may be a determinant of activity for namushens 3 and 4. These results confirm that namushen phorbol esters can mediate latency reversal when used in isolation from the parental crude extract.

### Namushen levels are broadly similar across multiple Mukungulu crude extracts but do not recapitulate all Mukungulu latency reversal activity *in vitro*

To determine relative abundance of namushens in crude Mukungulu, we next assessed namushen content level using LC-MS analysis (**Fig. 2**). For this analysis, we investigated both the crude Mukungulu extract described above (Extract A) as well as an independently prepared methanolic extract from Mukungulu collected from a separate location in a different year (Extract B) ^16^. This analysis determined that namushens made up 3.1% of total components in Extract A, comprising approximately equal proportions of namushens 1, 2, and 3 and approximately 70% less of namushens 4 and 5 (**Fig. 2A**; **Table S2**). In Extract B, namushens comprised 1.2% of total components, which consisted predominantly of namushen 2 followed by 3 and 1 (**Fig. 2A**). Linear regression analysis indicated good correlation of respective namushen levels between extracts (r^2^ = 0.59; **Fig. 2B**). These results indicate that namushens make up ∼1 to 3% of total soluble components of Mukungulu with proportions and amounts that are broadly consistent across independently sourced samples.

**Fig. 2.**
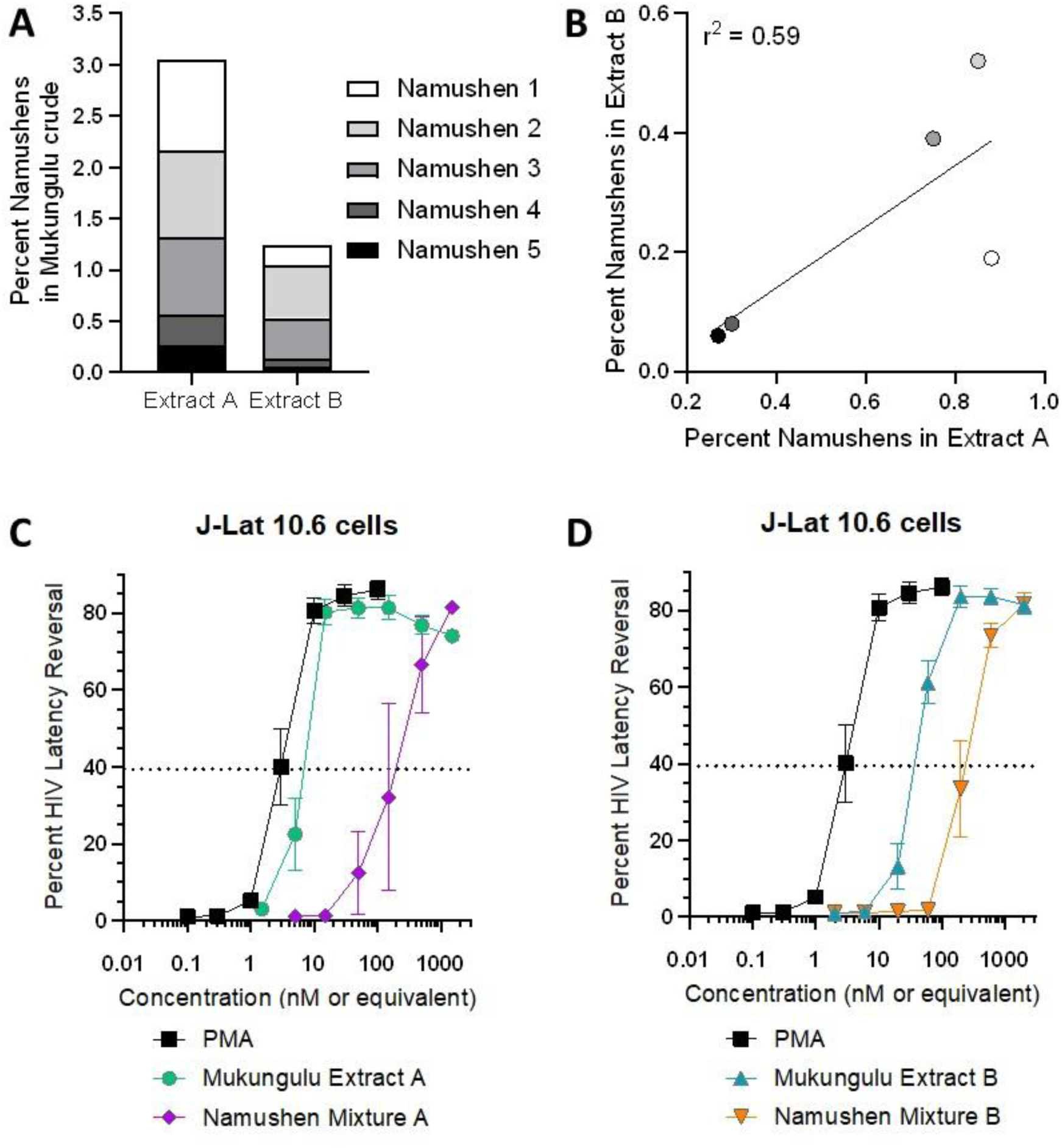
Namushen levels are broadly similar across different collections of Mukungulu medicinal plant. **A,** Steady-state polar metabolite analysis for namushens phytoconstituents in different Mukungulu crude extracts. **B,** Correlation of namushen levels in Mukungulu Extracts A and B. **C-D,** Effects of Mukungulu Extract A **(C)** and Extract B **(D)** on HIV latency reversal in J-Lat 10.6 cells when compared to namushens 1 – 5 recapitulated in equivalent molar concentrations to parental extracts.

We next asked whether namushens 1-5 were sufficient to recapitulate the full LRA magnitudes observed in crude extracts. We therefore reconstituted the five namushens in the same percentages and concentrations so they equal the same percentage of namushens to total volume present in parental extracts (i.e., identical to levels seen in **Fig. 2A**; **Table S2**). Namushen mixtures were then tested side-by-side with Batch A or B for ability to reverse latency in J-Lat 10.6 cells. As shown in **Fig. 2C**, treatment of J-Lat 10.6 cells with positive control PMA reversed latency with an EC_50_ of 1.0 nM, while Mukungulu Extract A exhibited a relative EC_50_ of 2.7 nM (where 1 µg/mL of Mukungulu is calculated to contain 20.6 nM of namushens; **Table S2**). By contrast, Namushen mixture A had an EC_50_ of only 130 nM, requiring a 48-fold higher concentration to achieve the same activity as the EC_50_ of intact Extract A (**Fig. 2C**). Similarly, while Extract B had a relative EC_50_ of 20.3 nM (where 1 µg/mL of Mukungulu is calculated to contain 51.0 nM of namushens; **Table S2**), namushen mixture B had an EC_50_ of only 150 nM, requiring a 7.4-fold higher concentration to achieve similar activity as that observed in intact Extract B (**Fig. 2D**). These results indicate that namushens 1-5, when applied in equal concentrations and proportions to those observed in parental intact Mukungulu extracts, do not fully recapitulate the latency reversal activity of Mukungulu crude extracts on their own.

### Mukungulu robustly reverses HIV-1 latency in PBMC from PLWH stably suppressed on ART

To investigate whether latency reversal observed in J-Lat T cell lines extended to primary cells, we obtained PBMC from 10 ART-suppressed PLWH. **Table 1** shows baseline characteristics of study participants. Having first established percent CD4+ T-cells in PBMC from each donor (**Fig. 3A**), we next quantified levels of intact and defective proviruses within isolated CD4+ T-cells from each study participant using the Intact Provirus DNA Assay (IPDA; **Fig. 3B-C**) ^20^. These results indicated an average of 573 ± 163 total proviruses per million CD4+ T-cells, consisting of 80 ± 25 intact proviruses and 493 ± 193 total defective proviruses per million CD4+ T-cells (representing 14.0 ± 4.3 and 86.0 ± 33.7% of total proviruses, respectively).

**Table 1.** Baseline characteristics of study participants.

| Patient ID | Assessed in CD4+ T-cell studies? | Color and shape on graphs | Age | Sex / Gender | Race | Ethnicity | Date of HIV Diagnosis (mm/yyyy) | Date of starting ART (mm/yyyy) | Years of documented continual viral suppression | CD4 nadir recorded at first diagnosis | Curent ART regimen | CD4 count on day of collection | Viral load on day of collection |
| --- | --- | --- | --- | --- | --- | --- | --- | --- | --- | --- | --- | --- | --- |
| 1 | N | Green-yellow hexagon | 45 | Male | Not Hispanic | African-American | 12/1998 | 2004 | 11 | 77 | Triumeq | 573 | < 20 |
| 2 | N | Gray down-triangle | 35 | Male | Unknown | Unknown | 01/2009 | 06/2010 | 12 | 201 | Triumeq | 478 | < 20 |
| 3 | Y | Green circle | 42 | Male | Not Hispanic | African-American | unk/1998 | 12/2013 | 8 | Unk, >200 | Juluca | 826 | < 20 |
| 4 | N | Peach circle | 43 | Male | Not Hispanic | African-American | 05/2003 | 2003 to 2006 | 12 | 349 | Symtuza | 653 | 51 |
| 5 | N | Green up-triangle | 45 | Male | Not Hispanic | African-American | 01/2011 | 2011 | 10 | 878 | Dovato | 531 | < 20 |
| 6 | Y | Orange diamond | 59 | Male | Not Hispanic | African-American | unk/1988 | 2004 | 10 | Unk, <20 | Biktarvy | 998 | < 20 |
| 7 | Y | Red down-triangle | 35 | Male | Not Hispanic | African-American | 2008 or 2009 | 2013 to 2014 | 2 | Unk | Stribild | 942 | < 20 |
| 8 | Y | Blue square | 60 | Male | Not Hispanic | White | unk/1998 | 06/1905 | 23 | Unk, >200 | Biktarvy | 457 | < 20 |
| 9 | N | Pink up-triangle | 51 | Male | Not Hispanic | White | 12/1997 | 1997 | 7 | 23 | Triumeq | 712 | < 20 |
| 10 | Y | Purple hexagon | 57 | Male | Not Hispanic | African-American | 01/2002 | 2004 | 17 | 425 | Biktarvy | 694 | < 20 |

**Fig. 3.**
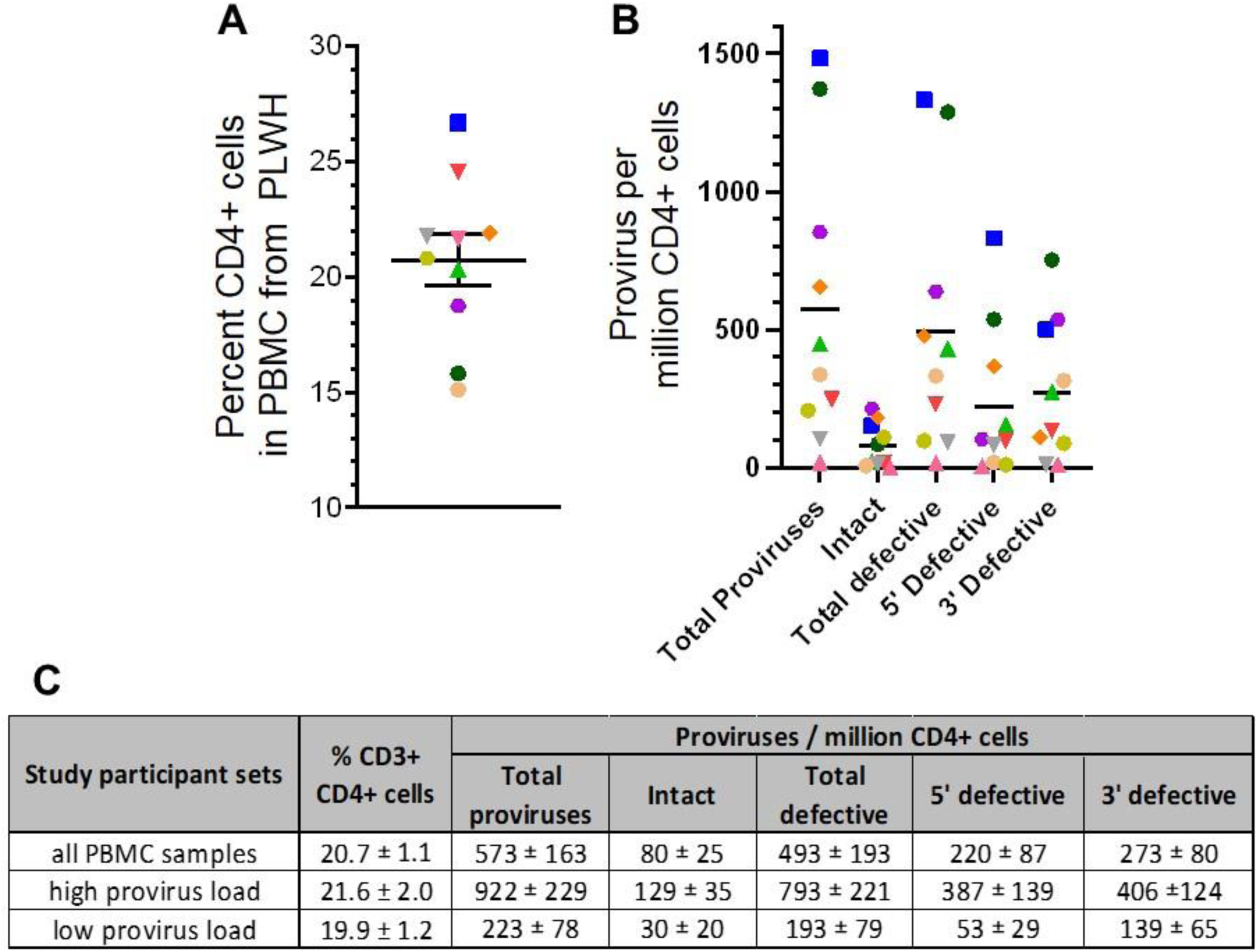
Baseline properties of primary cell samples obtained from 10 ART-suppressed PLWH. **A,** Percent CD3+CD4+ T-cells in PBMC. **B,** Provirus levels in isolated CD4+ T-cells from each donor, as measured by IPDA. In **A** and **B**, shapes and colors denote samples obtained from study participants as annotated in **Table 1**. **C,** Numerical summaries of data presented in **A-B**.

20 million PBMC from each of the 10 study participants were then cultured in triplicate in the presence of 1 µg/mL of Mukungulu (Extract B). As a positive control, PBMC were also treated in parallel with 50 µg/mL anti-CD3/CD28 dynabeads. After 72 hours of treatment, live cells were quantified by trypan blue stain. Using this approach, we found that PBMC treated with Mukungulu had 92.4 ± 3.5% viability relative to untreated PBMC (**Fig. 4A**), indicating no major cytotoxic effects that, at minimum, exceed cell replacement through LRA-induced proliferation.

**Fig. 4.**
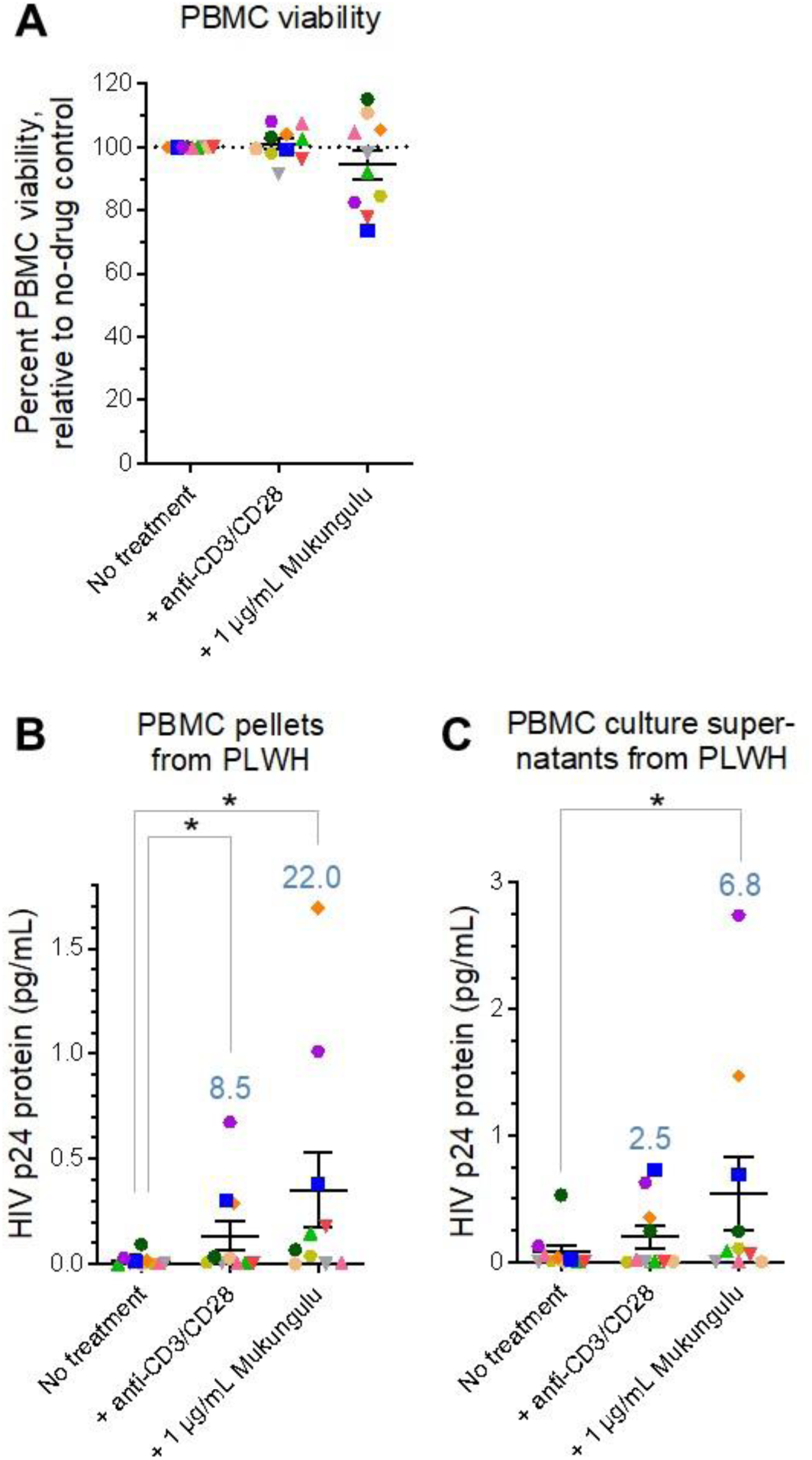
Mukungulu reactivates HIV expression in PBMC obtained from 10 ART-suppressed PLWH. **A,** Percent cell viability in the presence of LRAs after 72 hours treatment. Results are presented relative to viability of untreated cells cultured in parallel. **B-C**, Detection of gag-p24 protein in cell pellets (**B**) and culture supernatants (**C**) after 72 hours of treatment with LRAs, as measured by Simoa. In all panels, blue values denote average fold-increases over no-treatment controls. *, p < 0.05 as measured by one-sided Mann-Whitney test.

Cell pellets and culture supernatants were then collected from all experiments and assessed for viral protein production after 72 hours using HIV gag-p24 antigen detection by Simoa ^21^. In untreated cells, minimal HIV gag-p24 protein was detected across all cell pellets (average 0.016 ± 0.009 pg/mL), while anti-CD3/CD28 induced an average 0.136 ± 0.071 pg/mL of viral protein, or a significant 8.5-fold increase over no-drug control (p = 0.02; **Fig. 4B**). By contrast, Mukungulu treatment induced more gag-p24 production than anti-CD3/CD28 in almost all donors, resulting in an average 0.353 ± 0.178 pg/mL of viral protein. While this increase in viral protein was only 2.6-fold more than anti-CD3/CD28 induction (p = 0.2), it was 22.0-fold more than untreated controls (p = 0.03; **Fig. 4B**), indicating more robust latency reversal by Mukungulu than anti-CD3/CD28 following 72 hours of treatment. Similar results were observed from culture supernatants: while anti-CD3/CD28 treatment induced an average 0.199 ± 0.281 pg/mL of gag-p24 (a 2.5-fold increase over untreated cells with 0.079 ± 0.164 pg/mL of viral protein; p = 0.3), Mukungulu induced an average 0.542 ± 0.901 pg/mL of gag-p24. While this increase by Mukungulu was only 2.7-fold over anti-CD3/CD28 (p = 0.2), it produced a 6.8-fold increase over untreated cells (p = 0.04; **Fig. 4C**). These results suggest that Mukungulu also induces more latency reversal than anti-CD3/CD28 treatment in PBMC.

### Latency reversal induced by LRAs in PBMC isolated from PLWH correlates with intact HIV provirus levels in CD4+ T-cells

To explore whether LRA-induced viral protein production correlated with proviral load in CD4+ T-cells, we next compared HIV gag-p24 protein production by stimulated PBMC to provirus levels in isolated CD4+ T-cells as obtained by IPDA. As expected, viral protein expression did not correlate with percent CD3+ CD4+ T-cells, as the cohort had similar frequencies of CD4+ across all study participants (**Fig. 3A**). However, we observed that LRA-induced HIV-1 gag-p24 levels from both PBMC pellets and supernatants from cultured cells treated with anti-CD3/CD28 or Mukungulu correlated significantly with intact but not total or defective provirus levels (**Fig. 5**). For example, while limited correlation was observed for viral protein levels in PBMC pellets compared to either total provirus levels in CD4+ T-cells (r^2^ = 0.23; p = 0.16; **Fig. 5A**) or total defective provirus levels (r^2^ = 0.15; p = 0.27; **Fig. 5B**), a significant correlation was observed when compared to intact provirus levels (r^2^ = 0.74; p = 0.001; **Fig. 5C**). Similar correlations were also observed for viral protein from PBMC pellets treated with Mukungulu, with no correlation to total CD4+ T-cell provirus (r^2^ = 0.08; p = 0.44; **Fig. 5D**) or total defective provirus (r^2^ = 0.03; p = 0.63; **Fig. 5E**) but significant correlation with intact provirus levels (r^2^ = 0.60; p = 0.008; **Fig. 5F**). When culture supernatants were assessed, we similarly observed that HIV gag-p24 protein levels from both anti-CD3/CD28-treated and Mukungulu-treated culture supernatants significantly associated with intact provirus levels in CD4+ T-cells (anti-CD3/CD28 treatment: r^2^ = 0.71, p = 0.002; Mukungulu treatment: r^2^ = 0.74, p = 0.001; **Fig. 5G**). Taken together, results support that the magnitude of HIV reactivation induced by both anti-CD3/CD28 and Mukungulu in PBMC *ex vivo* associates with intact provirus levels in CD4+ T-cells.

**Fig. 5.**
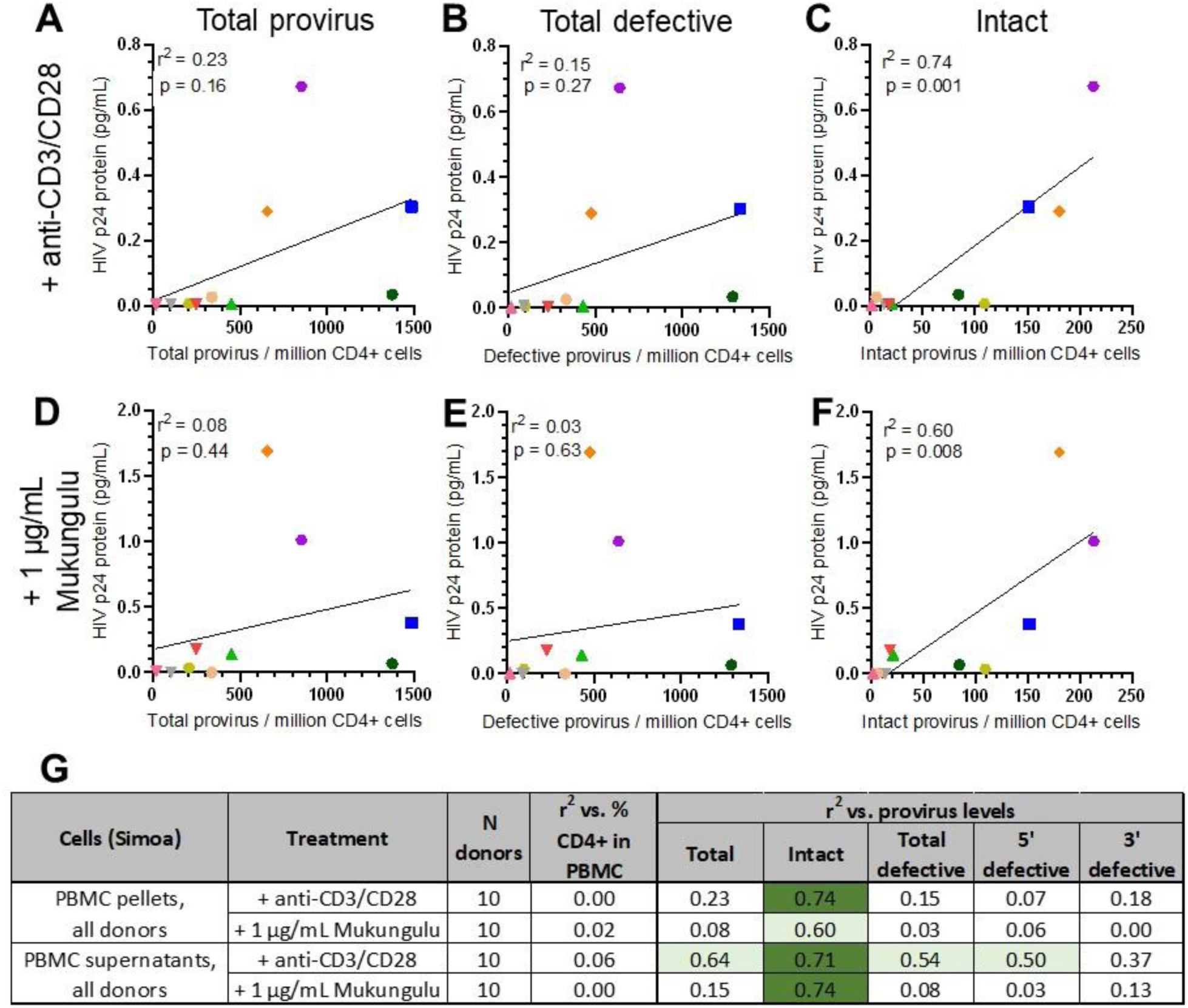
Magnitude of Mukungulu reactivation in PBMC correlates with intact provirus levels in CD4+ T-cells. **A-F**, Correlations of viral protein levels PBMC pellets following 72 hours treatment with anti-CD3/CD28 (**A-C**) or 1 µg/mL Mukungulu (**D-F**), as measured by simoa, relative to total provirus (**A, D**), total defective provirus (**B, E**) and intact proviruses (**C, F**) in isolated CD4+ T-cells. In all panels, colors/shapes denote individual donors as annotated in **Table 1**. **G,** Correlations of viral protein production relative to provirus levels in isolated CD4+ T-cells obtained from 10 PLWH. In panel **G**, light green shading denotes p < 0.05, and dark green shading denotes p < 0.005, as measured by ANOVA.

### Mukungulu reverses HIV-1 latency in CD4+ T-cells isolated from PLWH stably suppressed on ART

To reproduce and extend observations of latency reversal by Mukungulu in ART-suppressed PBMC to the context of CD4+ T-cells, we selected PBMC from study participants in the upper half of HIV provirus load (“high provirus load”), defined as having at least 240 total provirus copies per million CD4+ T-cells. This subset contained an average of 922 ± 229 total proviruses per million CD4+ T-cells, consisting of 129 ± 35 intact proviruses and 793 ± 221 total defective proviruses per million CD4+ T-cells (representing 14.0 ± 3.7 and 86.0 ± 24.0% of total proviruses, respectively; **Fig. 3C**).

CD4+ T-cells were next isolated from PBMC from each of the “high provirus load” subset, and 2 million CD4+ T-cells per donor were cultured in duplicate in the presence of 50 µg/mL anti-CD3/CD28 dynabeads or 1 µg/mL Mukungulu (Extract B). After 72 hours of treatment, cells in all conditions were again assessed for viability by trypan blue stain, where we found that CD4+ T-cells treated with Mukungulu had 94.4 ± 1.0% viability relative to untreated CD4+ T-cells tested in parallel, again indicating that it did not cause obvious cellular toxicity exceeding cell proliferation *ex vivo* (**Fig. 6A**). As expected, anti-CD3/CD28 treatment also did not cause obvious toxicity (98.1 ± 0.9% viability relative to untreated cells, respectively; **Fig. 6A**).

**Fig. 6.**
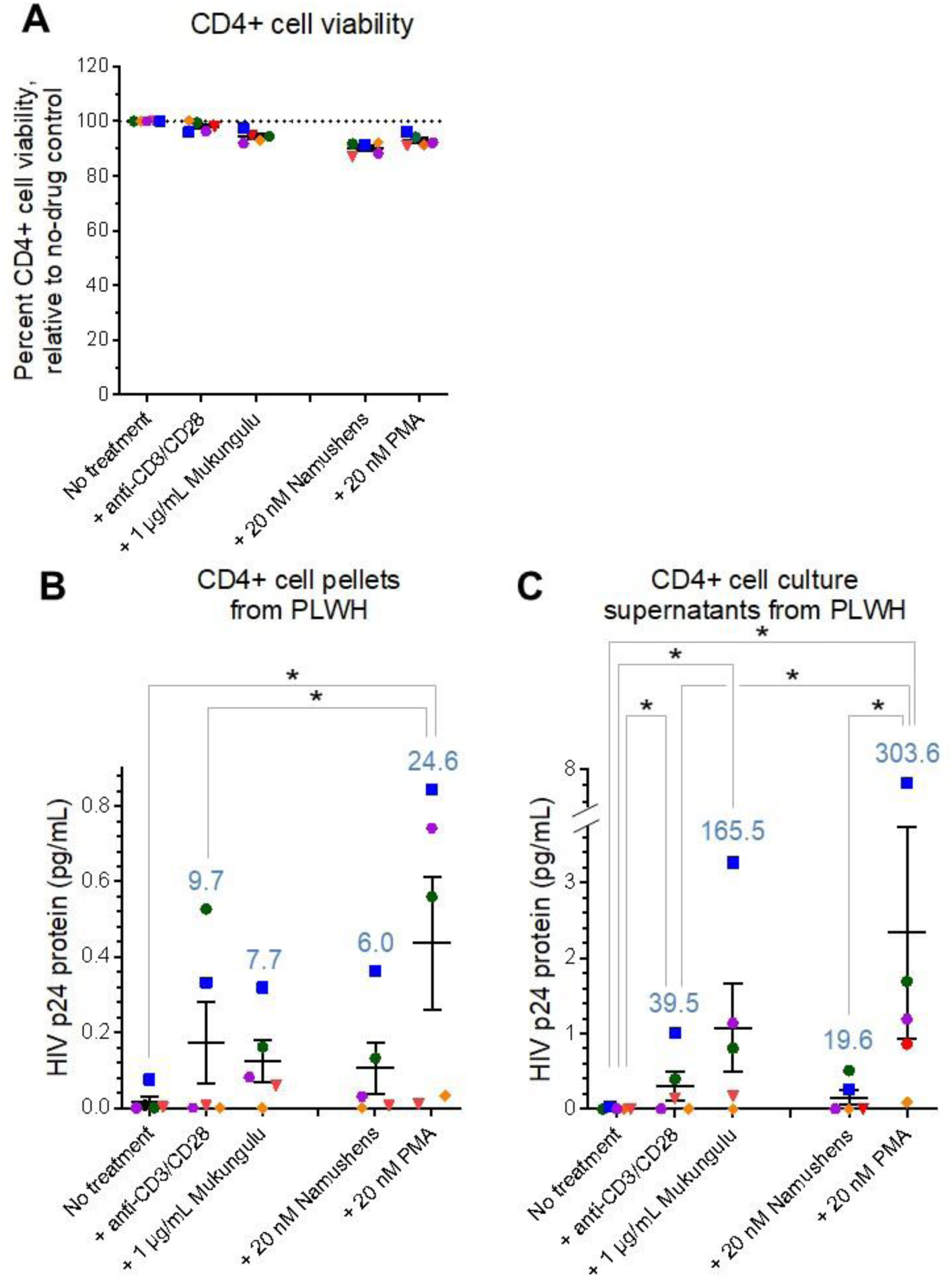
Mukungulu reactivates HIV expression in CD4+ T-cells obtained from 5 ART-suppressed PLWH. **A,** Percent cell viability in the presence of LRAs after 72 hours treatment. Results are presented relative to viability of untreated cells cultured in parallel. **B-C**, Detection of gag-p24 protein in cell pellets (**B**) and culture supernatants (**C**) after 72 hours of treatment with LRAs, as measured by Simoa. In all panels, blue values denote average fold-increases over no-treatment controls. *, p < 0.05 as measured by one-sided Mann-Whitney test.

Cell pellets and culture supernatants were again collected from all experiments and assessed by HIV gag-p24 Simoa. In untreated cells, minimal HIV gag-p24 protein was again detected across all CD4+ T-cell pellets (average 0.015 ± 0.012 pg/mL), while anti-CD3/CD28 induced an average 0.145 ± 0.094 pg/mL of viral protein. While this corresponded to an average 9.7-fold increase over untreated cells in viral protein production at 72 hours post-treatment, this increase was not statistically significant (p = 0.4; **Fig. 6B**). While Mukungulu also induced latency reversal in isolated CD4+ T-cells, with an average 0.114 ± 0.05 pg/mL of protein observed over 5 CD4+ T-cell samples, this corresponded to a 7.7-fold increase over untreated cells at 72 hours with borderline significance (p = 0.09; **Fig. 6B**). When culture supernatants were analyzed, we again found minimal gag-p24 protein produced from untreated cells (average 0.0006 ± 0.006 pg/mL). Anti-CD3/CD28 treatment induced an average 0.221 ± 0.154 pg/mL of viral protein in supernatants, corresponding to a significant 39.5-fold increase over untreated cells (p = 0.04; **Fig. 6C**). By contrast, Mukungulu induced 0.927 ± 0.467 pg/mL of viral protein, which corresponded to a 4.2-fold increase over anti-CD3/CD28-treated cells (p = 0.2) but a 165.5-fold increase over untreated cells (p = 0.03), similar to trends observed in PBMC.

To confirm whether the activity differences between Mukungulu and namushen mixtures (**Fig. 2C-D**) extended to *ex vivo* observations, we also treated in parallel CD4+ T-cells from 5 donors with 20 nM namushens (i.e., corresponding to concentration present in 1 µg/mL of Mukungulu; **Table S2**). As positive controls, isolated CD4+ T-cells were also treated with or 20 nM PMA, which represented maximal stimulation of J-Lat cells (as shown in **Fig. 2C-D)**. Following trypan blue stain, we found that CD4+ T-cells treated with namushens had 90.2 ± 1.1% viability relative to untreated CD4+ T-cells, while cells treated with PMA had 94.4 ± 1.0% viability (**Fig. 6A**). However, similar to *in vitro* observations, treatment with the corresponding namushen mixture on CD4+ T-cells did not show a significant increase in LRA activity in both cell pellets (6.0-fold; p = 0.2; **Fig. 6B**) and culture supernatants (19.6-fold; p = 0.1; **Fig. 6C**), whereas PMA achieved a significant induction in both (24.6-fold and 303.6-fold; p = 0.02 and 0.006, respectively; **Fig. 6B-C**).

Taken together, results demonstrate that Mukungulu is a robust latency reversing agent with direct activity on CD4+ T-cells with activity that is comparable to or exceeding that of anti-CD3/CD28 stimulation after 72 hours, particularly with more viral protein production than anti-CD3/CD28 treatment in culture supernatants. These data also support that an isolated namushen mixture is unlikely to match the viral antigen reactivation observed with Mukungulu, further indicating other factors in Mukungulu that may support HIV reactivation and replication over namushen-only based strategies.

### Mukungulu extract is tolerated and reverses latency reversal *in vivo*

To determine whether the amount of phorbol esters in Mukungulu would be tolerated by mice, we first injected a total of 18 CD-1 mice intraperitoneally with Mukungulu at 0.125, 1.25 or 12.5 mg/kg (3 female + 3 male mice per concentration) and monitored them for tolerance up to 72 hours post-administration. While no changes in behavior were observed for any mouse at 0.125 mg/kg administration, at higher concentrations mice began to exhibit hunched postures, slightly labored breathing, and reduced activity starting at 10 minutes post-administration which resolved after 8 hours. No other changes were observed including changes in body weight up to 72 hours post-administration, indicating that mice tolerated Mukungulu treatment at concentrations up to 12.5 mg/kg, albeit with temporary side effects.

We next investigated whether the latency reversal induced by Mukungulu *ex vivo* corresponded to *in vivo* efficacy using the ART-suppressed HIV-infected BLT humanized mice model ^23–26^. Here, 12 HIV-infected, ART-suppressed BLT humanized mice were placed into two experimental groups per the schematic study design shown in **Fig. 7A**. The test group comprised 7 mice receiving 5 mg/kg Mukungulu extract (Extract B), while the remaining 5 received PBS vehicle control (**Fig. 7A**). After 24 hours, mice were euthanized, and blood was collected to measure plasma viral load (pVL) and cellular viral RNA from isolated human CD4+ T-cells. Unlike CD-1 mice, BLT humanized mice treated with 5 mg/kg Mukungulu exhibited weight loss after 24 hours, where body weight was 93.5 ± 7.7% of weight before treatment, in contrast to PBS-treated mice where body weight was 99.8 ± 1.3% of pre-treatment (p = 0.05). This indicated poorer tolerance of Mukungulu by HIV-infected BLT humanized mice when compared to wild-type mice, which likely reflects experimental and/or strain-specific features. However, as shown in **Fig. 7B**, virus expression was readily detected following Mukungulu treatment, where an average pVL of 550 ± 130 viral copies/mL was observed, compared to the control cohort where the plasma viral load remained below the limit of detection (p = 0.009). Furthermore, and consistent with the plasma viral load rebound results, cellular viral RNA expression in isolated human CD4+ T-cells was observed in Mukungulu-treated mice averaging 1870 ± 250 viral mRNA copies per million cells compared to no detection in those animals that received vehicle control PBS (p = 0.03; **Fig. 7C**). Taken together, these results indicate that Mukungulu is a robust LRA at concentrations that can be tolerated *in vivo*.

**Fig. 7.**
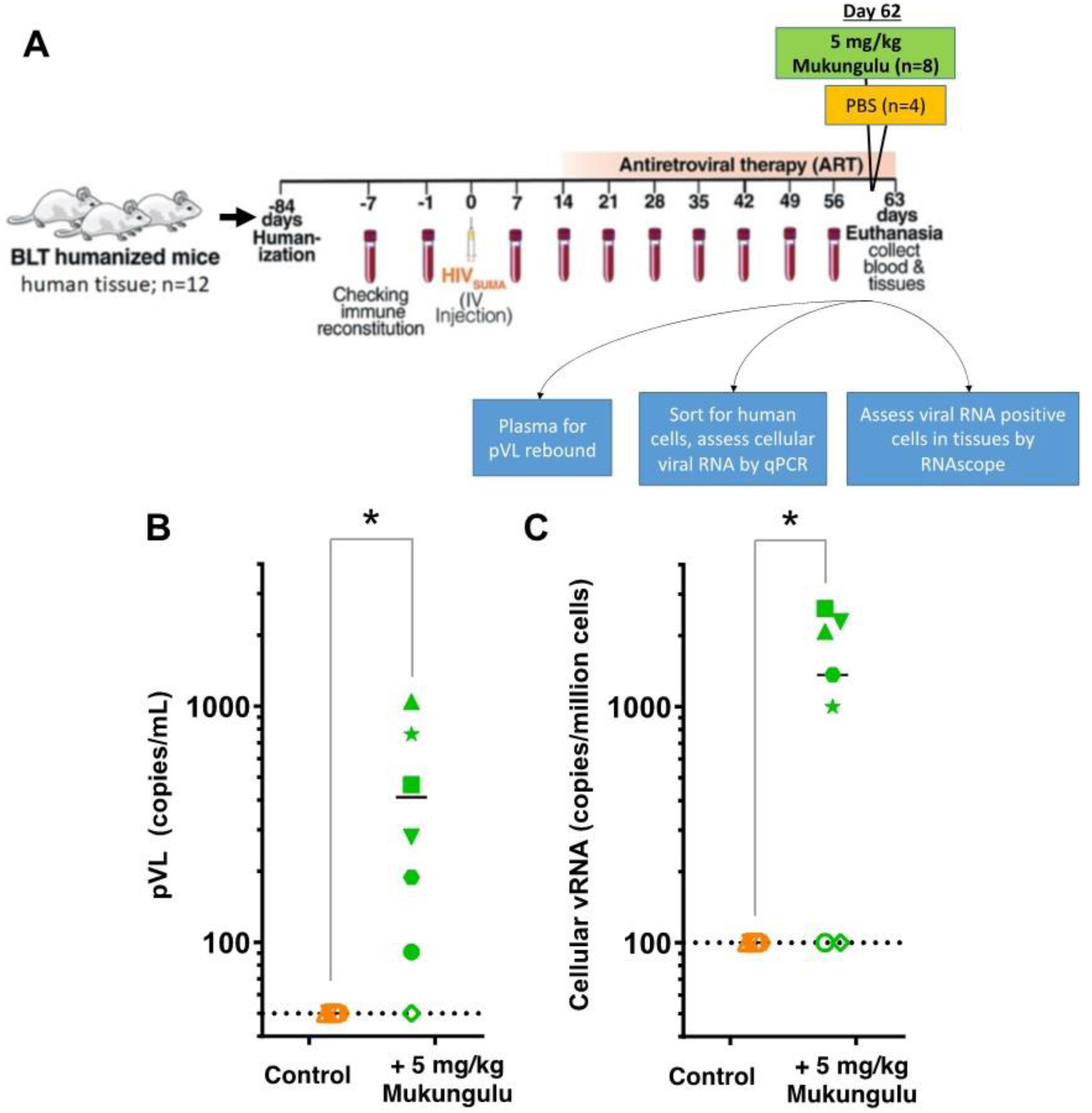
Mukungulu reverses HIV-1 latency *in vivo*. **A,** Schematic illustration of the BLT humanized mice model-based study (adapted and modified from ^25^). **B,** Plasma viral load (pVL) rebound measurements in ART-suppressed BLT humanized mice, treated with Mukungulu. **C,** Viral RNA induction from isolated human CD4+ T-cells from ART-suppressed mice treated with PBS vehicle control or Mukungulu. Colors/shapes denote individual mice. *, p < 0.05 as measured by one-sided Mann-Whitney test.

## Discussion

Having previously documented the traditional use of “Mukungulu,” a bark extract of *Croton megalobotrys*, to manage HIV/AIDS to supplement ART in Northern Botswana ^14,16^, here we show that Mukungulu can act as a robust LRA *ex vivo* in both PBMC and isolated CD4+ T-cells from PLWH as well as *in vivo* using a humanized mouse model. Mukungulu extract is also shown to be tolerated in uninfected control mice at doses able to induce latency reversal by an active component of Mukungulu-specific phorbol esters (namushens 1-5). Notably, the tolerability of Mukungulu stands in direct contrast to the established *in vivo* toxicities of PMA and other phorbol esters ^11–12^. Furthermore, as Mukungulu’s traditional use as a single dose correlates with documented improvement in patient health ^14,16^, its activity as an LRA described here raises the hypothesis that it may also be tolerated if used as an LRA strategy in conjunction with ART.

Extensive bioassay-guided fractionation of 81.3 grams of Mukungulu crude extract identified a total of 5 namushen phorbol esters. These namushens comprised 1.2 to 3.1% of the total phytochemical composition of Mukungulu and were present in similar proportions across two independent collections of Mukungulu extracts. However, when namushen mixtures were reconstituted to the concentrations and proportions seen in parental extracts, they did not recapitulate the full latency reversing profiles associated with their respective crude extracts in both J-Lat cells and culture supernatants from isolated CD4+ T-cells from ART-suppressed PLWH. These results suggest that isolated namushen mixtures reverse latency more slowly and/or that parental Mukungulu extracts more effectively induces viral expression after reactivation. While bioassay-guided fractionation indicates that isolated namushens are likely the dominant LRA species in Mukungulu, our results do not exclude the presence of additional, unidentified phytoconstituents in Mukungulu that could enhance namushen-driven latency reversal. Mukungulu reversal of HIV latency was also observed in both PBMC and isolated CD4+ T-cells from up to 10 ART-suppressed PLWH.

Notably, the activity of 1 µg/mL Mukungulu was generally on par with, or superior to, anti-CD3/CD28 polyclonal T-cell positive control activity. As expected, the higher the intact proviral load of HIV in CD4 T-cells, the better the detection of reactivation by LRAs *ex vivo*. We also observed that, to a first approximation, Mukungulu’s activity in isolated CD4+ T-cell pellets, where gag-p24 increased 7.7-fold relative to untreated cells, was lower than results from PBMC pellets, where Mukungulu induced a 22.0-fold increase, suggesting that Mukungulu may amplify its effects over 72 hours with other cells types apart from direct action on CD4+ T-cells.

Importantly, we used ART-suppressed humanized BLT mice infected with HIV ^23–26^ to demonstrate that a single 5 mg/kg injection of Mukungulu robustly reactivates HIV expression in most animals after 24 hours, as measured by both pVL and cell-associated HIV RNA levels. The magnitude of latency reversal induced by 5 mg/kg Mukungulu after 24 hours approximates what is reported for plasma viral RNA reactivation in a similar ART-suppressed BLT humanized, HIV-infected mouse model treated with 3 mg/kg of the SMAC mimetic AZD5582 after 48 hours ^27^, further supporting robust Mukungulu activity *in vivo*. Mice received a single dose of Mukungulu as this mirrors the traditional use Mukungulu, which is taken once followed by healer observation for 48 – 72 hours ^14–16^. It should be stressed, however, that HIV-infected humanized mice did not tolerate Mukungulu as well as control wild-type CD-1 mice, as measured by weight loss. We interpret this to be due to humanized mice being more sensitive to activation as created from immunodeficient mice known to be less robust than wild-type mice. Similar weight loss was also observed in postnatal Wistar rats following treatment with 10-fold lower concentrations of PMA at 500 µg/kg ^12^. It is also of interest to note our previous field documentation where we learned that traditional health practitioners do not administer Mukungulu to persons that appear excessively weak ^16^. Future investigation in both murine and nonhuman primate models will need to better define *in vivo* toxicity thresholds as well as changes in proinflammatory cytokine responses associated with *in vivo* Mukungulu dosing.

One important limitation to these studies is that all study participants were male and originated from the northeast United States, a predominant region for HIV subtype B. As subtype C is predominant in Botswana and Southern Africa, *ex vivo* studies need to be extended to specimens from PLWH in these areas, particularly given recent reports that LRAs may have more activity against latent subtype C proviruses relative to subtype B ^22^. Another limitation is the difference as to how Mukungulu was administered in mice when compared to traditional use in humans. Specifically, mice received an *i.p.* injection of crude Mukungulu extract, while humans are documented to ingest one leveled teaspoon of Mukungulu powder prepared in a small teacup filled with boiling hot porridge that is mixed until cold ^16^. Future studies will need to further investigate formulation strategies for Mukungulu administration relative to HIV reactivation activity.

In summary, we show that Mukungulu is a robust candidate LRA in primary cells from PLWH and in HIV humanized mouse models where reactivation activity doses were also documented to be tolerated despite the presence of PKC-activating phorbol esters. Identification of Mukungulu as an LRA strategy as a consequence of reverse pharmacology in collaboration with indigenous traditional medicine also stresses that current uses of medicinal plants globally may hold information on tolerability and impact on HIV disease yet to be identified.

## Acknowledgements

We are first and foremost indebted to the study participants who provided PBMC for this study. Funding was provided by the Canadian Institutes for Health Research (CIHR PJT-153057) (I.T.), the W.W. Smith Charitable Trust (A2301), the New Frontiers in Research Fund – Explorations (NFRFE-2018-01386) (I.T.), the Penn Center for AIDS Research Grant P30 AI 045008 (L.J.M and I.T.) and the Sub-Saharan African Network for TB/HIV Research Excellence (SANTHE) (K.R. and I.T.), a DELTAs African Initiative (grant no. DEL-15-006). The DELTAS Africa Initiative is an independent funding scheme of the African Academy of Sciences (AAS)’s Alliance for Accelerating Excellence in Science in Africa (AESA) and supported by the New Partnership for Africa’s Development Planning and Coordinating Agency (NEPAD Agency) with funding from the Wellcome Trust (grant no. 107752/Z/15/Z) and the United Kingdom government. G.M.L. was funded by NIAID NIH U24 AI143502. H.Y.T. was supported by NCI R50 CA221838. This work was also supported by the following grants to L.J.M.: Beyond Antiretroviral Treatment (BEAT)-HIV Delaney Collaboratory Grant UM1 AI164570, by the Robert I. Jacobs Fund of the Philadelphia Foundation, and the Herbert Kean, M.D., Family Professorship. K.R. was a recipient of a Canadian Queen Elizabeth II Diamond Jubilee Scholarship, a partnership between the Rideau Hall Foundation, Community Foundations of Canada, and Universities Canada, in addition to a SANTHE Ph.D. fellowship. Support for the Wistar Proteomics and Metabolomics Shared Resource was provided by NIH Cancer Center Support Grant CA010815, and the Thermo Scientific Q Exactive HF-X mass spectrometer was purchased with NIH grant S10 OD023586.

## Author contributions

Conceptualization: K.A.M., I.T., L.J.M. Methodology: K.R., Z.Y., H.Y.T., A.R.G., D.E.W., C.C., G.M.L., I.T. Validation: K.R., Z.Y., D.E.W., C.C. Formal Analysis: K.R., Z.Y., D.E.W., C.C., G.M.L., K.A.M., I.T. Investigation: K.R., Z.Y., H.Y.T., A.R.G., R.K., B.P., E.T.R., P.S., B.N.R., J.M., D.E.W., C.C., G.W., G.M.L. Resources: Z.Y., H.Y.T., K.M., G.M.L., P.Z., R.J.A., S.S., K.A.M., L.J.M. Writing – Original Draft: K.R., D.E.W., I.T. Writing – Review & Editing: I.T., L.J.M. Supervision: G.M.L., P.Z., R.J.A. Funding Acquisition: K.A.M., I.T., L.J.M.

## Materials and Methods

### Cells, viruses, animals, and reagents

J-Lat T cell clones 9.2 and 10.6 were obtained from the NIH AIDS Reagent Program, Division of AIDS, NIAID, NIH (contributed by Dr. Eric Verdin) ^18^. Cells were cultured in R10+ medium (RPMI 1640 with HEPES and L-glutamine, 10% fetal bovine serum, and 100 units/mL of penicillin, and 100 µg/mL streptomycin.

PBMCs and CD4+ T-cells were isolated from whole blood obtained from 10 ART-suppressed PLWH enrolled in the BEAT-HIV Collaboratory study (**Table 1**) by way of a written informed consent. At the time of blood samples collection, study participants had plasma HIV RNA levels below limit of detection (< 20 copies/mL plasma viral load; only one donor had 51 copies/mL pVL). Study protocols and ethics were approved and granted by Institutional Review Boards at The Wistar Institute (Philadelphia, USA) and Philadelphia FIGHT, as guided by the US Department of Health and Human Services guidelines pertaining the human subject research.

Study protocols were approved by the Institutional Animal Care and Use Committee of the Wistar Institute and adhere to the *NIH Guide for the Care and Use of Laboratory Animals*.

### Collection of Mukungulu

*Croton megalobotrys* Müll Arg. (“Mukungulu”) bark was collected by the traditional healer SS in 2018 around Maun, Ngamiland District, North-Western Botswana. Four hundred and eighty grams (480 g) of the bark was grinded and extracted with CH_2_Cl_2_/methanol as previously described^16^ giving rise to 102.8 g of an oily dark brown extract (Yield: 21.4%). 81.3 g of this extract served as the starting material for the bioassay-guided fractionation reported here (Extract A). Extract B represented the methanolic extract from original *Croton megalobotrys* bark collected by the healer SS in 2014 in the Kazungula District covering the Zambia/Botswana border region, which was botanically authenticated as previously described ^16^ (Voucher specimen KM-Ks-3-2015). Ethical approval and subsequent research permits were granted by the Ministry of Health, Botswana (Permit no.: PPME:13/18/1Vol VIII (354)) and and the Ministry of Infrastructure, Science and Technology, Botswana (Permit no: ETH 5 (1), respectively.

### In vitro HIV-1 latency reversal model

HIV-1 latency reversal profiles of test agents (Mukungulu Extracts and its novel derivatives) were measured as previously described ^17^ using flow cytometry-based green fluorescent protein (GFP) reporter cells (J-Lat 10.6 cells) that contain an integrated HIV-1 provirus, modelling HIV-1 latency *in vitro* ^18^. Basically, J-Lat 10.6 cells were plated in 96-well plates (i.e., 2 × 10^5^ cells per well at a final volume of 200 µL) and treated with multiple concentrations of test agents, alongside an established positive control LRA, PMA and incubated at 37 °C and 5% CO_2_ for 24 hours. Cells were then assessed and detected for GFP-expression using flow cytometry, where GFP-positive cells indicated latent HIV-1 reactivation. 0.1% DMSO was used as a vehicle control.

### Bioassay-guided fractionation of Mukungulu extracts

Latency reversal properties of Mukungulu extract fractions were determined as described above using J-Lat 9.2 cells. Details on the isolation and structural characterization of namushens 1 – 5 can be found in the **Supplementary Materials**.

### Namushen quantitation in Mukungulu extracts

Mukungulu extracts were each diluted 20-, 40- and 80-fold with 80% methanol. Diluted samples were analyzed by LC-MS using a Thermo Scientific Q Exactive HF-X mass spectrometer in-line with a Thermo Scientific Vanquish UHPLC system and reversed-phase Synergi Polar-RP C18 column (Phenomenex). Replicate injections were performed for each sample. namushens were detected as formate adducts in negative ion mode, similar to published results for related phorbol esters ^28^. The concentrations of namushens were determined from MS peak integrations using calibration curves generated from purified compounds diluted into 80% methanol (0.005 – 10 μM range, quadratic fit, 1/x^2^ weighting). Quantitation was highly reproducible with most namushens in the Mukungulu extracts having coefficients of variation of less than 10% among the replicate injections and across different dilutions.

### Percent CD3+ CD4+ T-cell detection in PBMCs

PBMC suspensions were stained with the following antibodies: CD4-V450 (clone: RPA-T4; BD Biosciences) and CD3-PE-CF594 (clone: UCHT1; BD Biosciences) as described by the manufacturer. Data were collected on a BD Biosciences LSRII flow cytometer.

### Intact Provirus DNA Assay (IPDA) quantification of persistent HIV-1 proviral DNA

In this study, we employed the IPDA method to quantify persistent intact and defective HIV-1 proviral DNA in CD4+ T-cells isolated from 10 study participants. An in-depth description of the IPDA design rationale is available in Bruner, *et al.* ^20^. Sample processing and IPDA measurements were performed by Accelevir Diagnostics under company standard operating procedures by blinded operators. Briefly, cryopreserved PBMCs were viably thawed, and total CD4+ T-cells were obtained via negative immunomagnetic selection (EasySep Human CD4+ T-cell Enrichment Kit, Stemcell Technologies), with cell count, viability, and purity assessed by flow cytometry both before and after selection. Genomic DNA was isolated using the QIAamp DNA Mini Kit (Qiagen) with RNA removed by RNase A treatment. DNA concentrations were determined by fluorometry (Qubit dsDNA BR Assay Kit, Thermo Fisher Scientific), and DNA quality was determined by ultraviolet-visible (UV/VIS) spectrophotometry (QIAxpert, Qiagen). Genomic DNA was then analyzed by IPDA using the Bio-Rad QX200 AutoDG Droplet Digital PCR system, and results were reported as frequencies of intact, defective, and total proviruses per million input cells.

### HIV-1 gag-p24 Single-Molecule Assay (Simoa)

CD4+ T-cells were isolated from frozen PBMCs from 5 study participants using the EasySep Human CD4+ T-Cell Enrichment Kit (Stemcell Technologies). 2 million CD4+ T-cells were then cultured in 96-well plates in duplicate in 200 μL R10+ media supplemented with 100 U/mL IL-2, 200 nM Raltegravir, and appropriate test agents. For PBMC-based studies, 20 million cells were cultured in 12-well plates in triplicate in 2 mL of R10+ plus IL-2, raltegravir, and test agents. Cells were then incubated for 72 hours at 37 °C and 5% CO_2_. Following incubation, live cells were quantified by trypan blue stain, and cell pellets and culture supernatants were harvested for HIV gag-p24 protein Simoa. Cell pellets were resuspended in Simoa buffer (49% Blocker^TM^ Casein in PBS (Thermo-Fisher), 49% FBS, 1% triton, 1x protease inhibitor cocktail) and incubated for 30 minutes, while culture supernatants were mixed with 10% triton (in PBS) to a final concentration of 1%, before storage at −80 °C.

Samples were then analyzed for gag-p24 protein using Simoa methods described previously by Wu et al., 2021 ^21^. For detection of viral protein, anti-p24 conjugated magnetic beads were added to thawed cell pellet lysate in Simoa buffer and supernatant samples to enrich for gag-p24. Bound gag-p24 was eluted with 100 μL 0.1% trifluoroacetic acid (TFA), and the eluate was neutralized with 20 µL of 1M Tris-Hcl at pH 9.0. Meanshile, the beads were eluted with 100 μL of 3% BSA in PBS again to obtain residual p24. Both eluates were combined into a 200 μL final volume. 140 μL of each sample was then analyzed on a Quanterix HDX-1 platform using gag-p24 Simoa kits obtained from Quanterix. Gag-p24 concentrations were calculated with Simoa software that uses four-parameter logistic regression curve fitting and reported as pg/mL of cell lysate or culture supernatant.

### in vivo tolerability of Mukungulu

Tolerability studies were performed at Alliance Pharma (Malvern, PA, USA). 18 mice (9 female + 9 male) were injected intraperitoneally with 0.125, 1.25, or 12.5 mg/kg of Mukungulu crude extract (n = 6 each) and evaluated changes in behavior for the first 30 minutes post-administration, every 4-6 hours for the first 24 hours, and then every 12-24 hours up to 72 hours.

### Generation of bone marrow-liver-thymus (BLT) humanized mice

Two independent cohorts of BLT mice were generated as previously described ^23–26^ in accordance with the Wistar Institute Animal Care and Research Committee regulations (protocol 201360). Briefly, female NSG mice aged 6-8 weeks (NOD.Cg-Prkdcscid Il2rgtm1Wjl/SzJ, Jackson Laboratory) were pretreated with busulfan at 30mg/kg and implanted with human fetal thymic tissue fragments and fetal liver tissue fragments under the murine renal capsule. Following surgery, mice were injected via the tail vein with CD34+ hematopoietic stem cells. Human fetal liver and thymus tissues were procured from Advanced Bioscience Resources (Alameda, CA). Twelve weeks post-surgery, human immune cell reconstitution in peripheral blood was determined by flow cytometry.

### HIV infection, antiretroviral therapy suppression, and Mukungulu treatment

BLT mice from each cohort were randomly divided into two groups and infected intravenously with 10^4^ x 50% tissue culture infectious dose (TCID_50_) of HIV_SUMA_. Peripheral blood was collected weekly for plasma viral load assays. Two weeks after infection, mice were placed on a diet combined with ART (1500 mg/kg Emtricitabine, 1560 mg/kg Tenofovir-Disoproxil-Fumarate, and 600 mg/kg Raltegravir). Five weeks post-ART, mice were treated with phosphate-buffered saline (PBS) control or 5 mg/kg Mukungulu by intraperitoneal injection. After 24 hours, mice were euthanized, and blood was collected. Plasma viral loads were measured as previously described ^23–26^. Single-cell suspensions were generated using the gentleMACS Octo Dissociator (San Diego, CA). RNA was extracted using AllPrep DNA/RNA/ miRNA Universal Kit (Qiagen, catalog # 80224). Cell associated HIV RNA was measured as previously described ^23–26^.

### Data and statistical analyses

All results were analyzed using Graphpad Prism v. 10.2.0 (Boston, MA, USA). For *in vitro* studies, results denote the mean ± s.e.m. from at least 3 independent experiments. For *ex vivo* studies, each data point denotes average results from a given study participant sample performed in duplicate (for CD4+ T-cells) or triplicate (for PBMC). For *in vivo* studies, each data point denotes average results from a given animal performed in duplicate. Statistical significance was determined using a one-sided Mann-Whitney test or ANOVA, where a *P* value < 0.05 was considered significant.

## References

1. T. W. Chun, L. Stuyver, S. B. Mizell, L.A. Ehler, J. A. Mican, M. Baseler, A. L. Lloyd, M. A. Nowak, A. A. Fauci. Presence of an inducible HIV-1 latent reservoir during highly active antiretroviral therapy. Proc. Natl. Acad. Sci. USA 94, 13193–13197 (1997).

2. D. Finzi, J. Blankson, J., J. D. Siliciano, J. B. Margolick, K. Chadwick, T. Pierson, K. Smith, J. Lisziewicz, F. Lori, C. Flexner, T. C. Quinn, R. E. Chaisson, E. Rosenberg, B. Walker, S. Gange, J. Gallant, J., R. F. Siliciano. Latent infection of CD4+ T cells provides a mechanism for lifelong persistence of HIV-1, even in patients on effective combination therapy. Nat. Med. 5, 512–517 (1999).

3. A. Chawla, C. Wang, C. Patton, M. Murray, Y. Punekar, A. de Ruiter, C. Steinhart. A Review of Long-Term Toxicity of Antiretroviral Treatment Regimens and Implications for an Aging Population. Inf. Dis. Ther. 7, 183–195 (2018).

4. S. Zicari, L. Sessa, N. Cotugno, A. Ruggiero, E. Morrocchi, C. Concato, S. Rocca, P. Zangari, E. C. Manno, P. Palma. Immune Activation, Inflammation, and Non-AIDS Co-Morbidities in HIV-Infected Patients under Long-Term ART. Viruses. 11, 200 (2019).

5. M. Massanella, R. Fromentin, N. Chomont. Residual inflammation and viral reservoirs: Alliance against an HIV cure. Curr. Opin. HIV AIDS. 11, 234–241 (2016).

6. S. G. Deeks. HIV: Shock and kill. Nature. 487, 439–440 (2012).

7. I. Sadowski, F. B. Hashemi. Strategies to eradicate HIV from infected patients: elimination of latent provirus reservoirs. Cell. Mol. Life Sci. 76, 3583–3600 (2019).

8. U. Mbonye, J. Karn. The cell biology of HIV-1 latency and rebound. Retrovirology. 21, 6 (2024).

9. J. M. Zerbato, H. V. Purves, S. R. Lewin, T. A. Rasmussen. Between a shock and a hard place: challenges and developments in HIV latency reversal. Curr. Opin. Virol. 38, 1–9 (2019).

10. Q. Debrabander, K. S. Hensley, C. K. Psomas, W. Bramer, T. Mahmoudi, B. J. van Welzen, A. Verbon, C. Rokx. The efficacy and tolerability of latency-reversing agents in reactivating the HIV-1 reservoir in clinical studies: A systematic review. J. Virus Erad. 9, 100342 (2023).

11. S. E. Smith, B. S. Meldrum. The protein kinase C activators, phorbol 12-myristate,13-acetate and phorbol 12,13-dibutyrate, are convulsant in the pico-nanomolar range in mice. Eur. J. Pharmacol. 213, 133–135 (1992).

12. M. Dzietko, M. Hahnemann, O. Polley, M. Sifringer, U. Felderhoff-Mueser, C. Bührer. Effects of PMA (Phorbol-12-myristate-13-acetate) on the developing rodent brain. Biomed. Res. Int. 2015, 318306 (2015).

13. B. Patwardhan, A. D. B. Vaidya. Natural products drug discovery: accelerating the clinical candidate development using reverse pharmacology approaches. Indian J. Exp. Biol. 48, 220–227 (2010).

14. I. Tietjen, B. N. Ngwenya, G. Fotso, D. E. Williams, S. Simonambango, B. T. Ngadjui, R. J. Andersen, M. A. Brockman, Z. L. Brumme, Z. L., K. Andrae-Marobela. The Croton megalobotrys Müll Arg. traditional medicine in HIV/AIDS management: Documentation of patient use, in vitro activation of latent HIV-1 provirus, and isolation of active phorbol esters. J. Ethnopharmacol. 211, 267–277 (2018).

15. B. Tembeni, A. Sciorillo, L. Invernizzi, T. Klimkait, L. Urda, P. Moyo, D. Naidoo-Maharaj, N. Levitties, K. Gyampoh, G. Zu, Z. Yuan, K. Mounzer, S. Nkabinde, M. Nkabinde, N. Gqaleni, I. Tietjen, L. J. Montaner, V. Maharaj. HPLC-Based Purification and Isolation of Potent Anti-HIV and Latency Reversing Daphnane Diterpenes from the Medicinal Plant *Gnidia sericocephala* (*Thymelaeaceae*). Viruses. 14, 1437 (2022).

16. I. Tietjen, T. Gatonye, B. N. Ngwenya, A. Namushe, S. Simonambanga, M. Muzila, P. Mwimanzi, J. Xiao, D. Fedida, Z. L. Brumme, M. A. Brockman, K. Andrae-Marobela. Croton megalobotrys Müll Arg. and Vitex doniana (Sweet): Traditional medicinal plants in a three-step treatment regimen that inhibit in vitro replication of HIV-1. J. Ethnopharmacol. 191, 331–340 (2016).

17. K., Richard, C., Schonhofer, L. B. Giron, J. Rivera-Ortiz, S. Read, T. Kannan, N. N. Kinloch, A. Shahid, R. Feilcke, S. Wappler, P. Imming, M. Harris, Z. L. Brumme, M. A. Brockman, K., Mounzer, A. V. Kossenkov, M. Abdel-Mohsen, K. Andrae-Marobela, L. J. Montaner, I. Tietjen. The African natural product knipholone anthrone and its analogue anthralin (dithranol) enhance HIV-1 latency reversal. J. Biol. Chem. 295, 14084–14099 (2020).

18. A. Jordan, D. Bisgrove, E. Verdin. HIV reproducibly establishes a latent infection after acute infection of T cells in vitro. EMBO J. 22, 1868–1877 (2003).

19. P. Hashemi, K. Barreto, W. Bernhard, A. Lomness, N. Honson, T. A. Pfeifer, P. R. Harrigan, I. Sadowski. Compounds producing an effective combinatorial regimen for disruption of HIV-1 latency. EMBO Mol. Med. 10, 160–174 (2018).

20. K. M. Bruner, Z. Wang, F. R. Simonetti, A. M. Bender, K. J. Kwon, S. Sengupta, E. J. Fray, S. A. Beg, A. R. R. Antar, K. M. Jenike, L. N. Bertagnolli, A. A. Capoferri, J. T. Kufera, A. Timmons, C. Nobles, J. Gregg, N. Wada, Y. C. Ho, H. Zhang, J. B. Margolick, J. N. Blankson, S. G. Deeks, F. D. Bushman, J. D. Siliciano, G. M. Laird, R. F. Siliciano. A quantitative approach for measuring the reservoir of latent HIV-1 proviruses. Nature. 566, 120–125 (2019).

21. G. Wu, C. Cheney, Q. Huang, D. J. Hazuda, B. J. Howell, P. Zuck. Improved detection of HIV gag p24 protein using a combined immunoprecipitation and digital ELISA method. Front. Microbiol. 12, 636703 (2021).

22. U. Ranga, A. Panchapakesan, C. Saini. HIV-1 subtypes and latent reservoirs. Curr. Opin. HIV AIDS. 19, 87–92 (2024).

23. Z. Yuan, G. Kang, L. Daharsh, W. Fan, Q. Li. SIVcpz closely related to the ancestral HIV-1 is less or non-pathogenic to humans in a hu-BLT mouse model. Emerg. Microb. Inf. 7, 59 (2018).

24. Z. Yuan, G. Kang, F. Ma, W. Lu, W. Fan, C. M. Fennessey, B. F. Keele, Q. Li. Recapitulating Cross-Species Transmission of Simian Immunodeficiency Virus SIVcpz to Humans by Using Humanized BLT Mice. J. Virol. 90, 7728–7739 (2016).

25. Z. Yuan, L. B. Giron, C. Hart, K. Gyampoh, J. Koshy, K. Y. Hong, T. Niki, T. A. Premeaux, L. C. Ndhlovu, C. Deleage, L. J. Montaner, M. Abdel-Mohsen. Human galectin-9 promotes the expansion of HIV reservoirs in vivo in humanized mice. AIDS. 37, 571–577 (2023).

26. N.L. Board, Z. Yuan, F. Wu, M. Moskovljevic, M. Ravi, S. Sengupta, S. S. Mun, F. R. Simonetti, J. Lai, P. Tebas, K. Lynn, R. Hoh, S. G. Deeks, J. D. Siliciano, L. J. Montaner, R. F. Siliciano. Bispecific antibodies promote natural killer cell-mediated elimination of HIV-1 reservoir cells. Nat. Immunol. 25, 462–470 (2024).

27. C. C. Nixon, M. Mavigner, G. C. Sampey, A. D. Brooks, R. A. Spagnuolo, D. M. Irlbeck, C. Mattingly, P. T. Ho, N. Schoof, C. G. Cammon, G. K. Tharp, M. Kanke, Z. Wang, R. A. Cleary, A. A. Upadhyay, C. De, S. R. Wills, S. D. Falcinelli, C. Galardi, H. Walum, N. J. Schramm, J. Deutsch, J. D. Lifson, C. M. Fennessey, B. F. Keele, S. Jean, S. Maguire, B. Liao, E. P. Browne, R. G. Ferris, J. H. Brehm, D. Favre, T. H. Vanderford, S. E. Bosinger, C. D. Jones, J. P. Routy, N. M. Archin, D. M. Margolis, A. Wahl, R. M. Dunham, G. Silvestri, A. Chahroudi, J. V. Garcia. Systemic HIV and SIV latency reversal via non-canonical NF-κB signalling in vivo. Nature. 578, 160–165 (2020).

28. M. P. Neu, S. Schober, M. Mittelbach. Quantification of Phorbol Esters in Jatropha curcas by HPLC-UV and HPLC-ToF-MS with Standard Addition Method. Eur. J. Lipid. Sci. Tech. 120, 1800146 (2018).

29. E. Abner, A. Jordan. HIV “shock and kill” therapy: In need of revision. Antivir. Res. 166, 19–34 (2019).

30. J. B. McBrien, M. Mavigner, L. Franchitti, S. A. Smith, E. White, G. K. Tharp, H. Walum, K. Busman-Sahay, C. R. Aguilera-Sandoval, W. O. Thayer, R. A. Spagnuolo, M. Kovarova, A. Wahl, B. Cervasi, D. M. Margolis, T. H. Vanderford, D. G. Carnathan, M. Paiardini, J. D. Lifson, J. H. Lee, J. T. Safrit, S. E. Bosinger, J. D. Estes, C. A. Derdeyn, J. V. Garcia, D. A. Kulpa, A. Chahroudi, G. Silvestri. Robust and persistent reactivation of SIV and HIV by N-803 and depletion of CD8+ cells. Nature. 578, 154–159 (2020).

31. S. Mutascio, T. Mota, L. Franchitti, A. A. Sharma, A. Willemse, S. N. Bergstresser, H. Wang, M. Statzu, G. K. Tharp, J. Weiler, R. P. Sékaly, S. E. Bosinger, M. Paiardini, G. Silvestri, R. B. Jones, D. A. Kulpa. CD8+ T cells promote HIV latency by remodeling CD4+ T cell metabolism to enhance their survival, quiescence, and stemness. Immunity. 56, 1132–1147 (2023).

