## Supplemental Info for "*Ex vivo* and *in vivo* HIV-1 latency reversal by “Mukungulu,” a protein kinase C-activating African medicinal plant extract"

Richard et al.

**Supplementary Materials**

1. Isolation and identification of novel compounds from the Mukungulu plant crude extract
2. Tables S1-S2
3. Figures S1-S6

**Isolation and identification of novel compounds from the Mukungulu plant crude extract.**

General experimental procedures, collection and extraction of plant material and isolation of namushens 1-5 were as described previously for the isolation of namushens 1 and 2 [37].

**Namushen-3** was isolated as a clear glass [UV (4:1 MeCN/$H_2O$) $\lambda_{max}$ 207, 253 nm; $^1H$ and $^{13}C$ NMR see **Table S1**; positive ion HRESITOFMS $[M + Na]^+$ *m/z* 623.3554 (calculated for $C_{35}H_{52}O_8Na$, 623.3554), appropriate for a molecular formula of $C_{35}H_{52}O_8$, requiring 10 sites of unsaturation, one more than that observed in namushens-1 and 2]. Comparison of the $^1H$/$^{13}C$/gCOSY/gHSQC/gHMBC/tROESY NMR data of namushen-3 **(Table S1, Figures S1-S2)** with that of namushens-1 and 2 along with the additional site of unsaturation revealed that in namushen-3 the 2-methylbutanoate ester residue at C-17 in namushen-1 had been replaced by the olefin containing tiglic acid residue and that namushen-3 could be assigned the structure drawn in **Figures S1-S2**.

**Namushen-4** was isolated as a clear glass [UV (9:1 MeCN/$H_2O$) $\lambda_{max}$ 201, 249 nm; $^1H$ and $^{13}C$ NMR see **Table S1**; positive ion HRESITOFMS $[M + Na]^+$ *m/z* 651.3884 (calculated for $C_{37}H_{56}O_8Na$, 651.3867), appropriate for a molecular formula of $C_{37}H_{56}O_8$, requiring 10 sites of unsaturation, the same as namushen-3]. The NMR spectra of namushen-3 and namushen-4 were remarkably similar **(Table S1, Figures S3-S4)**, and the structures of the two compounds were found to only differ in the length of the alkyl chain of the lipid of the ester functionality at C-13. The structure of namushen-4 was assigned as drawn in **Figures S3-S4**.

**Namushen-5** was isolated as a clear glass [UV (9:1 MeCN/$H_2O$) $\lambda_{max}$ 200, 235 nm; $^1$H and $^{13}$C NMR see **Table S1**; positive ion HRESITOFMS [M + Na]$^+$ *m/z* 525.3190 (calculated for $C_{30}H_{46}O_6Na$, 525.3187), appropriate for a molecular formula of $C_{30}H_{46}O_6$, requiring 8 sites of unsaturation]. Comparison of the NMR spectra of namushen-5 **(Table S1, Figures S5-S6)** to that of namushens-3 and 4 revealed that namushen-5 lacked substitution at C-17 and was assigned the structure drawn in **Figures S5-S6**.

**Table S1.** $^{13}C$ and $^{1}H$ NMR Data for Namushens 3-5 and Comparison with Namushen-1 Recorded in $C_6D_6$.

| | Namushen-3 | | Namushen-4 | | Namushen-5 | | Namushen-1 | |
|---|---|---|---|---|---|---|---|---|
| Position # | $\delta_C$ | $\delta_H$ (*J* in Hz) | $\delta_C$ | $\delta_H$ (*J* in Hz) | $\delta_C$ | $\delta_H$ (*J* in Hz) | $\delta_C$ | $\delta_H$ (*J* in Hz) |
| 1 | 160.4 | 7.42 (bs) | 160.6 | 7.43 (bs) | 160.9 | 7.48 (bs) | 160.4 | 7.42 (bs) |
| 2 | 133.0 | / | 133.0 | / | 132.9 | / | 132.9 | / |
| 3 | 208.1 | / | 208.3 | / | 208.7 | / | 208.1 | / |
| 4 | 73.8 | / | 73.8 | / | 74.0 | / | 73.8 | / |
| 5 | 38.8 | 2.23 (bd, 18.9)<br>2.36 (bd, 18.9) | 38.8 | 2.30 (bd, 18.9)<br>2.39 (bd, 18.9) | 38.6 | 2.43 (bd, 19.1)<br>2.48 (bd, 19.1) | 38.8 | 2.25 (bd, 18.9)<br>2.37 (bd, 18.9) |
| 6 | 140.6 | / | 140.7 | / | 140.0 | / | 140.6 | / |
| 7 | 129.0 | 5.61 (bd, 5.2) | 129.1 | 5.63 (bd, 5.0) | 129.3 | 5.71 (bd, 5.1) | 129.0 | 5.60 (bd, 5.4) |
| 8 | 38.9 | 3.13 (bt, 5.2) | 38.9 | 3.16 (bt, 5.0) | 38.9 | 3.17 (bt, 5.1) | 38.9 | 3.11 (bt, 5.4) |
| 9 | 76.1 | / | 76.2 | / | 75.6 | / | 76.1 | / |
| 10 | 56.2 | 3.43 (bs) | 56.2 | 3.45 (bs) | 55.6 | 3.48 (bm) | 56.2 | 3.43 (bs) |
| 11 | 36.6 | 2.05[b] | 36.6 | 2.08[b] | 36.1 | 2.13 (m) | 36.6 | 2.05[b] |
| 12 | 32.5 | 1.79 (m)<br>2.07[b] | 32.5 | 1.79 (dd, 16.7, 13.7)<br>2.08[b] | 31.9 | 1.79 (dd, 14.2, 11.3)<br>2.08[b] | 32.4 | 1.78 (dd, 16.5, 13.7)<br>2.05[b] |
| 13 | 63.5 | / | 63.5 | / | 63.1 | / | 63.5 | / |
| 14 | 31.5 | 1.21 (d, 5.2) | 31.5 | 1.22 (d, 5.0) | 31.5 | 0.96 (d, 5.1) | 31.6 | 1.20 (d, 5.4) |
| 15 | 26.8 | / | 26.8 | / | 22.2 | / | 26.9 | / |
| 16 | 11.5 | 1.19 (s) | 11.6 | 1.20 (s) | 14.9 | 1.04 (s) | 11.5 | 1.18 (s) |
| 17 | 69.7 | 4.24 (d, 11.4)<br>4.28 (d, 11.4) | 69.7 | 4.24 (d, 11.3)<br>4.29 (d, 11.3) | 22.8 | 1.18 (s) | 69.3 | 4.17 (d, 11.3)<br>4.21 (d, 11.3) |
| 18 | 18.9 | 0.96 (d, 6.1) | 18.9 | 0.97 (d, 6.6) | 18.3 | 1.00 (d, 6.3) | 18.8 | 0.96 (d, 5.9) |
| 19 | 10.0 | 1.58 (dd, 2.9, 1.2) | 10.0 | 1.58 (bd, 1.8) | 9.4 | 1.58 (dd, 2.6, 1.1) | 10.0 | 1.58 (dd, 2.9, 1.3) |
| 20 | 67.8 | 3.58 (dd, 12.8, 5.3)<br>3.63 (dd, 12.8, 5.3) | 67.9 | 3.62 (d, 12.9)<br>3.68 (d, 12.9) | 67.5 | 3.70 (d, 12.8)<br>3.76 (d, 12.8) | 67.8 | 3.59 (d, 12.8)<br>3.64 (d, 12.8) |
| 20-OH | - | 0.63 (t, 5.3) | | / | - | / | - | / |
| 1’ | 175.7 | / | 175.7 | / | 175.0 | / | 175.7 | / |
| 2’ | 34.6 | 2.07[b] | 34.6 | 2.08[b] | 34.1 | 2.07 (t, 7.6) | 34.6 | 2.10 (m) |
| 3’ | 25.0 | 1.48 (m) | 25.0 | 1.50 (m) | 24.5 | 1.51 (m) | 25.0 | 1.51 (m) |
| 4’ | 29.3 | 1.14[b] | 29.4 | 1.16[b] | 28.7 | 1.16[b] | 29.57 | 1.18[b] |
| 5’ | 29.58[a] | 1.14-1.23 | 30.03[a] | 1.16-1.32 | 29.10[a] | 1.16-1.22 | 29.36[a] | 1.16-1.27 |
| 6’ | 29.65[a] | 1.14-1.23 | 30.03[a] | 1.16-1.32 | 28.98[a] | 1.16-1.22 | 29.64[a] | 1.16-1.27 |
| 7’ | 29.74[a] | 1.14-1.23 | 29.82[a] | 1.16-1.32 | 28.90[a] | 1.16-1.22 | 29.74 | 1.16-1.27 |
| 8’ | 32.2 | 1.23[b] | 29.78[a] | 1.16-1.32 | 31.6 | 1.22[b] | 32.2 | 1.24[b] |
| 9’ | 23.1 | 1.30 (m) | 29.61[a] | 1.16-1.32 | 22.4 | 1.29 (m) | 23.1 | 1.30 (m) |
| 10’ | 14.0 | 0.91 (t, 7.0) | 32.3 | 1.28[b] | 13.7 | 0.91 (t, 7.1) | 14.3 | 0.91 (t, 7.3) |
| 11’ | - | - | 23.1 | 1.30 (m) | - | - | - | - |
| 12’ | - | - | 14.4 | 0.92 (t, 7.3) | - | - | - | - |
| 1” | 167.5 | / | 167.5 | / | - | - | 175.9 | / |
| 2” | 129.2 | / | 129.2 | / | - | - | 41.3 | 2.33 (qdd, 7.0, 7.0, 7.0) |
| 3” | 136.9 | 6.99 (qq, 7.2, 1.4) | 136.9 | 6.98 (qq, 7.1, 1.3) | - | - | 27.2 | 1.40 (dqd, 13.6, 7.0, 7.0)<br>1.72 (dqd, 13.6, 7.0, 7.0) |
| 4” | 14.0 | 1.38 (dq, 7.2, 1.0) | 14.0 | 1.38 (d, 7.1) | - | - | 11.8 | 0.88 (t, 7.0) |
| 5” | 12.2 | 1.84 (m) | 12.2 | 1.84 (bs) | - | - | 16.9 | 1.12 (d, 7.0) |

[a]Assignments within a column are interchangeable. [b]Multiplicity not determined due to overlapping signals/chemical shifts determined from 2D data.

**Table S2.** Determination of relative levels of namushens in crude Mukungulu batches A and B.

| **Namushens** | Mukungulu Extract A | | Mukungulu Extract B | |
|---|---|---|---|---|
| | Amount per 1 μg/mL | | Amount per 1 μg/mL | |
| | ng | nM | ng | nM |
| 1 | 1.9 | 3.2 | 8.8 | 14.6 |
| 2 | 5.2 | 8.3 | 8.5 | 13.7 |
| 3 | 3.9 | 6.5 | 7.5 | 12.4 |
| 4 | 0.8 | 1.6 | 3.0 | 5.9 |
| 5 | 0.6 | 0.9 | 2.7 | 4.2 |
| Total | 12.4 | 20.6 | 30.4 | 51.0 |
| Percent of Crude Extract | 1.2% | | 3.0% | |

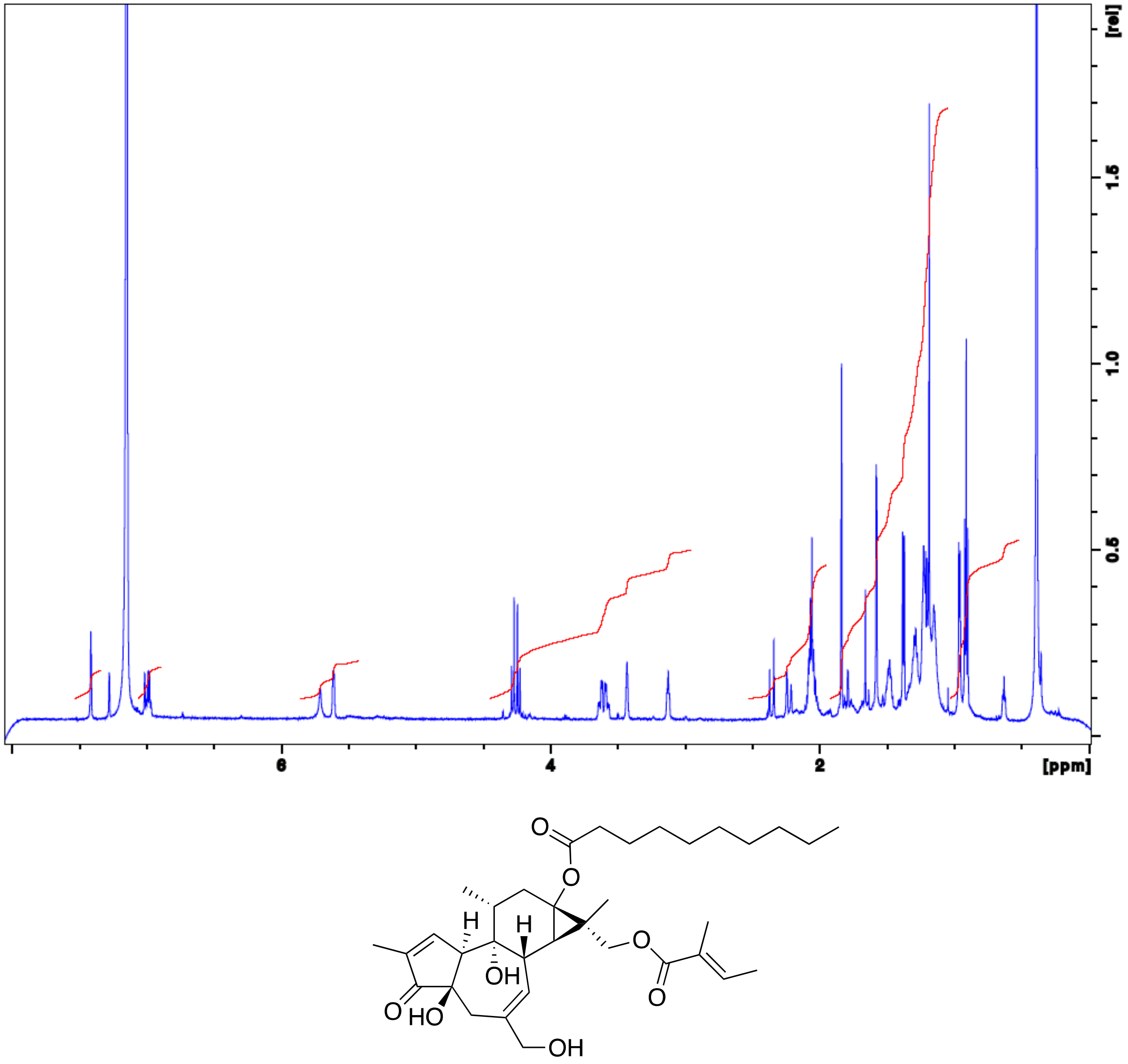


**Figure S1.** $^{1}H$ NMR Spectrum of namushen-3 recorded at 600 MHz in $C_6D_6$.

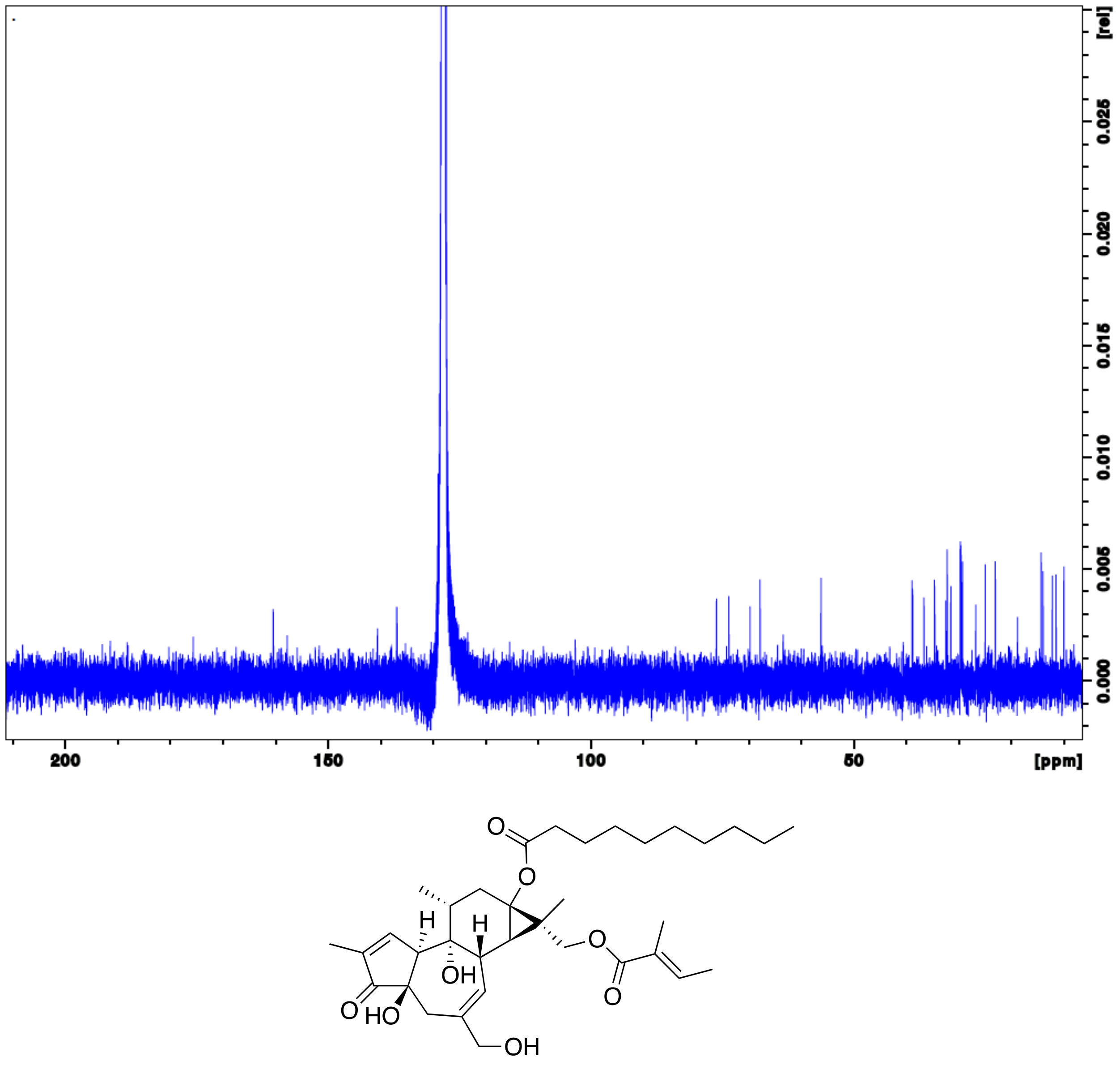


**Figure S2.** $^{13}C$ NMR Spectrum of namushen-3 recorded at 150 MHz in $C_6D_6$.

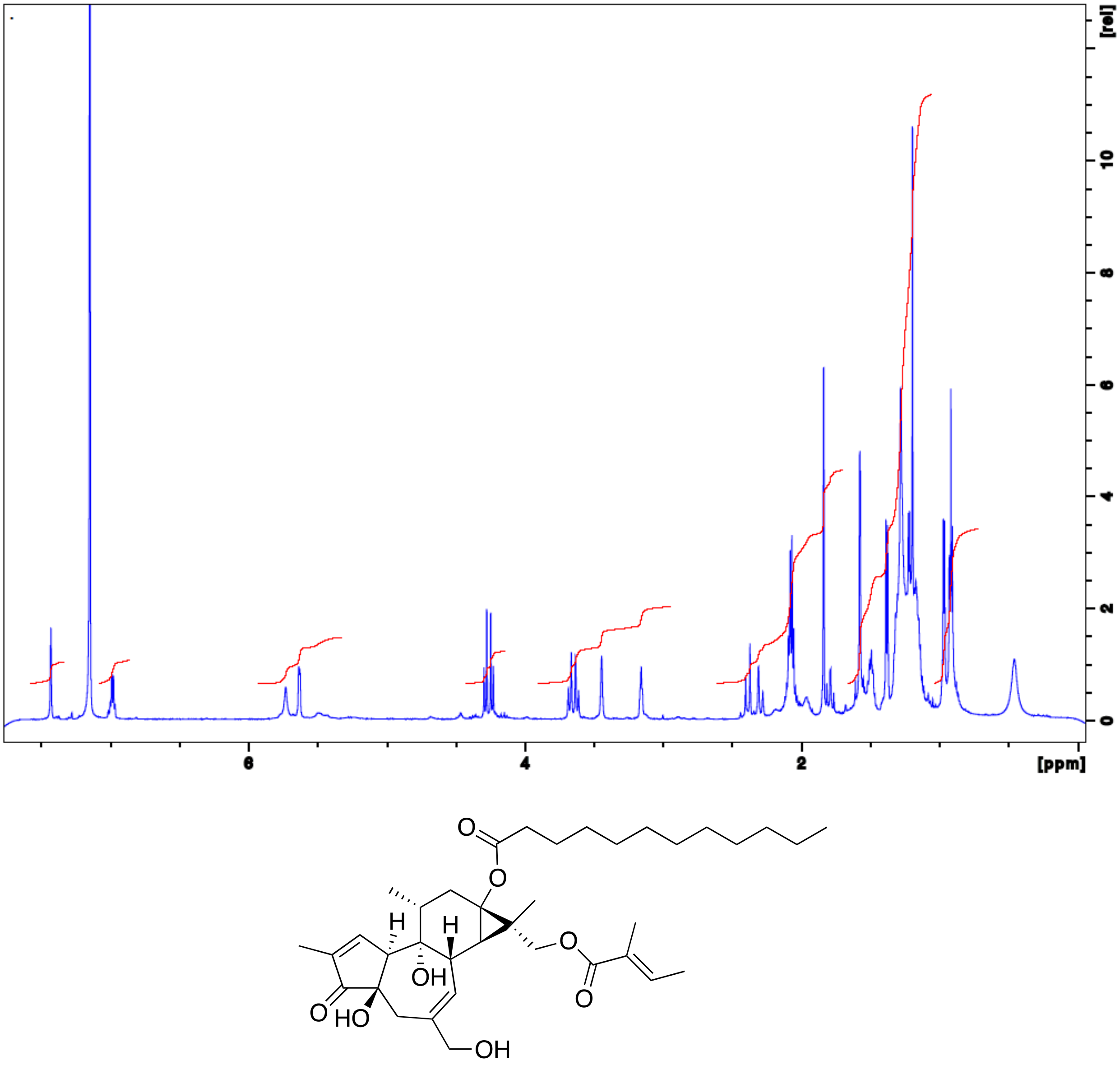


**Figure S3.** $^{1}H$ NMR Spectrum of namushen-4 recorded at 600 MHz in $C_6D_6$.

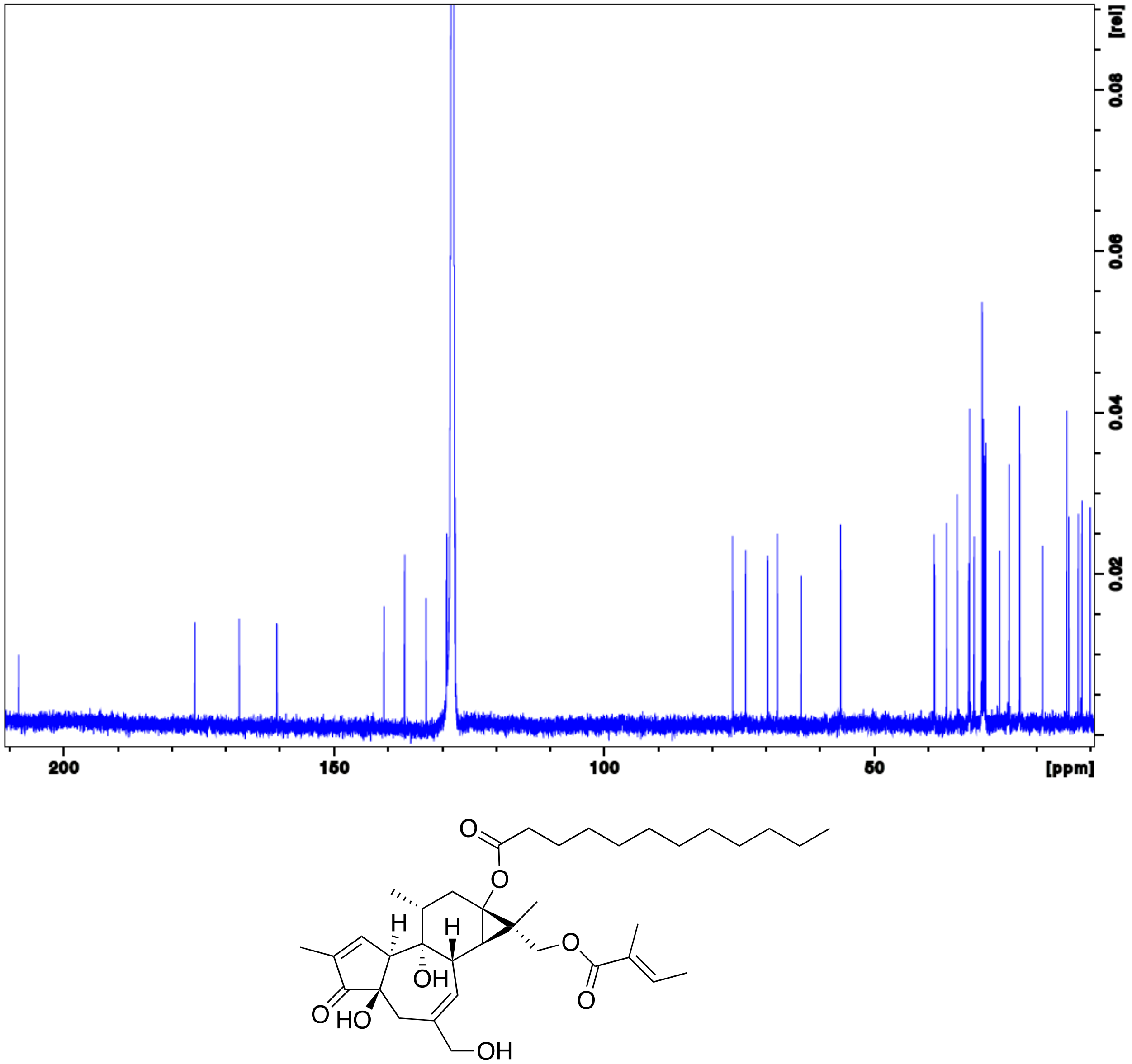


**Figure S4.** $^{13}C$ NMR Spectrum of namushen-4 recorded at 150 MHz in $C_6D_6$.

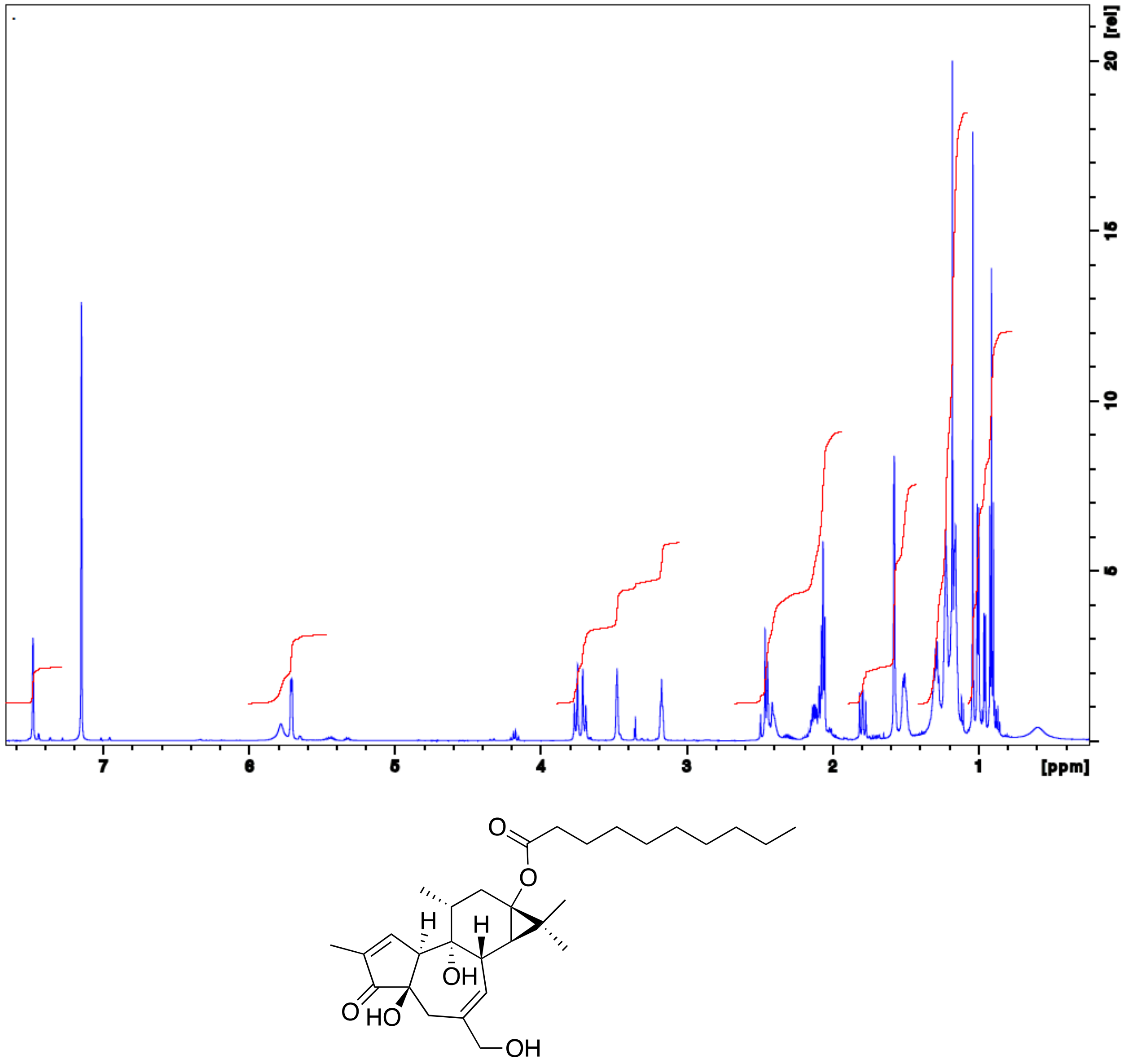


**Figure S5.** [1]H NMR Spectrum of namushen-5 recorded at 600 MHz in $C_6D_6$.

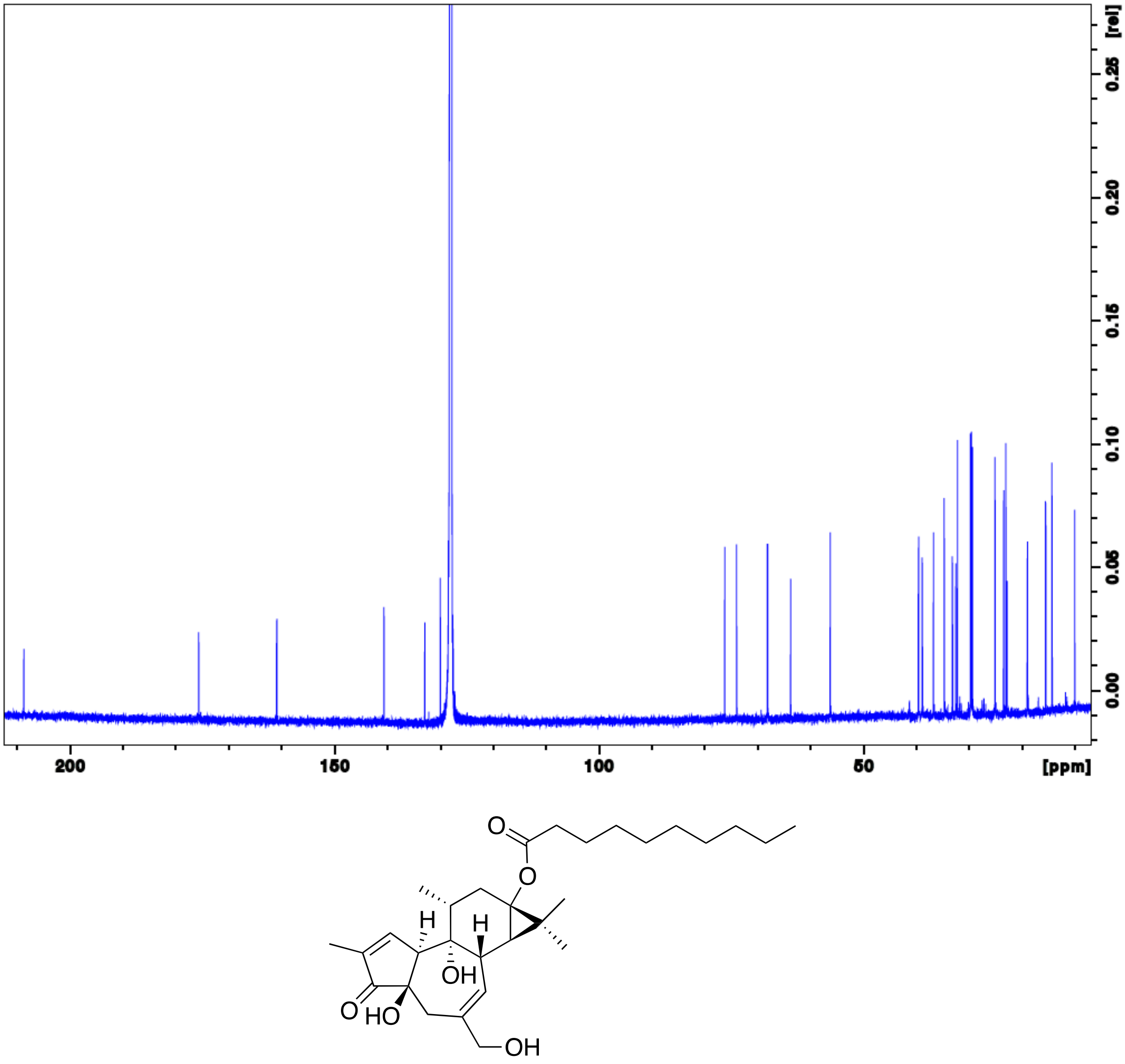


**Figure S6.** $^{13}C$ NMR Spectrum of namushen-5 recorded at 150 MHz in $C_6D_6$.
